# Generation of a Retina Reporter hiPSC Line to Label Progenitor, Ganglion, and Photoreceptor Cell Types

**DOI:** 10.1101/658963

**Authors:** Phuong T. Lam, Christian Gutierrez, Katia Del Rio-Tsonis, Michael L. Robinson

## Abstract

Early in mammalian eye development, *VSX2*, *BRN3b*, and *RCVRN* expression marks neural retina progenitors (NRPs), retinal ganglion cells (RGCs), and photoreceptors (PRs), respectively. The ability to create retinal organoids from human induced pluripotent stem cells (hiPSC) holds great potential for modeling both human retinal development and retinal disease. However, no methods allowing the simultaneous, real-time monitoring of multiple specific retinal cell types during development currently exist. Here, we describe a CRISPR/Cas9 gene editing strategy to generate a triple transgenic reporter hiPSC line (PGP1) that utilizes the endogenous *VSX2*, *BRN3b*, and *RCVRN* promoters to specifically express fluorescent proteins (Cerulean in NR<u>**P**</u>s, eGFP in R<u>**G**</u>Cs and mCherry in <u>**P**</u>Rs) without disrupting the function of the endogenous alleles. Retinal organoid formation from the PGP1 line demonstrated the ability of the edited cells to undergo normal retina development while exhibiting appropriate fluorescent protein expression consistent with the onset of NRPs, RGCs, and PRs. Organoids produced from the PGP1 line expressed transcripts consistent with the development of all major retinal cell types. The PGP1 line offers a powerful new tool to study retinal development, retinal reprogramming, and therapeutic drug screening.

## INTRODUCTION

Stem cells provide multiple avenues to address irreversible vision loss associated with retinal cell death caused by trauma or by diseases including Age-Related Macular Degeneration (AMD) and glaucoma. AMD and glaucoma alone leave millions of people blind and cost billions of dollars in social welfare and lost productivity^1,2^. Current treatments for these diseases may slow the progression of retinal degeneration, but no available treatments restore lost retinal tissue. Recent advancements in 3D-culture have led to the development of retinal organoids, that consist of all major retinal cell types from human induced pluripotent stem cells (hiPSCs)^3^. This capability means that these hiPSC-derived organoids can experimentally model normal human retina development as well as onset and progression of retinal disease. Additionally, these organoids can provide a platform to screen new drugs for the treatment or the cure of retinal disease^3,4^. Finally, hiPSCs provide the ability to make unlimited numbers of specific retinal neurons for potential transplantation therapies.

The rapidly expanding use of hiPSCs to model retinal development and disease justifies the need for tools to monitor the appearance of specific cell types without having to interrupt the normal developmental process. CRISPR/Cas9 gene editing greatly expands the ability to insert cell-type-specific reporter genes into hiPSCs for this purpose. Although several hiPSCs with single reporter knock-ins exist^5–7^, these lines only detect terminally differentiated retinal cell types. This limits the ability to study progression of retinal development in real time. In addition, having multiple cell type reporters inserted in the same genome will permit specific cell sorting and allow for the optimization of protocols that enrich for the development of one retinal cell type over another. For example, one might need a large number of photoreceptors or ganglion cells to screen for drugs to treat AMD or glaucoma, respectively^8,9^. To address these limitations, we utilized CRISPR/Cas9 genome editing to create a hiPSC retina reporter line to monitor the development of neural retina progenitor (NRP), retinal ganglion (RGC), and photoreceptor (PR) cells.

Several important considerations went into choosing a strategy to target NRPs, RGCs and PRs without damaging the ability of the engineered hiPSC line to undergo normal retinal development. *VSX2* encodes a transcription factor that marks NRPs before they differentiate into mature retinal cell types^10^. In the mature retina, only post-mitotic bipolar cells express *VSX2*^11–13^. RGCs represent the first mature retinal neuron cell type to develop and most RGCs express the *BRN3b* transcription factor^12^. PRs (rods and cones) express *RCVRN*, a calcium binding protein important in the recovery phase of visual excitation^14^. The cell-type-specific expression pattern of these genes made them appropriate endogenous targets for the insertion of fluorescent reporter genes. Since these three genes all play an important role in retina development and/or function, the ideal reporter hiPSC line will retain the function of both alleles of all three of these genes. Therefore, we utilized viral P2A peptides to create fusion genes between the target and the fluorescent reporter that would self-cleave upon translation^15^.

The value of a multiple-targeted retina reporter line depends on its ability to differentiate faithfully into all retinal cell types in a stereotypical fashion while simultaneously providing visible readout without the need to add dyes or to stop the developmental process. Here, we inserted a P2A:Cerulean reporter into the *VSX2* locus, a P2A:eGFP reporter into the *BRN3b* locus and a P2A:mCherry reporter into the *RCVRN* locus. In each case, we used CRISPR/Cas9 genome editing to replace the stop codon of the endogenous gene with the P2A reporter. Validation of the resultant triple transgenic hiPSC (PGP1) line came from the ability of these cells to generate retinal organoids with the appropriate expression of each fluorescent reporter gene.

## RESULTS

### Creation of the Neural Retina <u>P</u>rogenitor/ Retinal <u>G</u>anglion/ <u>P</u>hotoreceptor (PGP1) Reporter hiPSC Line by CRISPR/Cas9 Genome Editing

CRISPR/Cas9 genome editing technology provided a powerful platform to create a hiPSC line targeted with multiple fluorescent reporter genes driven by endogenous cell-type-specific promoters. In order to achieve cell-type-specific fluorescent protein expression without inactivating the targeted gene, we designed a strategy to replace the stop codon of endogenous genes with a sequence encoding a P2A peptide fused to a fluorescent reporter gene by homology directed repair (HDR, Fig. 1 A). To facilitate the insertion of multiple different reporter genes, the HDR targeting construct for each gene contained a different antibiotic resistance cassette. Cells nucleofected with both a CRISPR/Cas9 vector (containing the *S. pyogenes* Cas9 coding sequence and a sgRNA), and an HDR targeting construct were selected with the appropriate antibiotics. DNA sequence analysis of at least two PCR amplicons (see Table S1 for primer sequences) encompassing the 5’ and 3’ ends of the targeted modification with at least one PCR primer outside of the sequence contained in the homology arm (HA) confirmed each homologous recombination event (Fig. 1 B-D).

**Figure 1.**
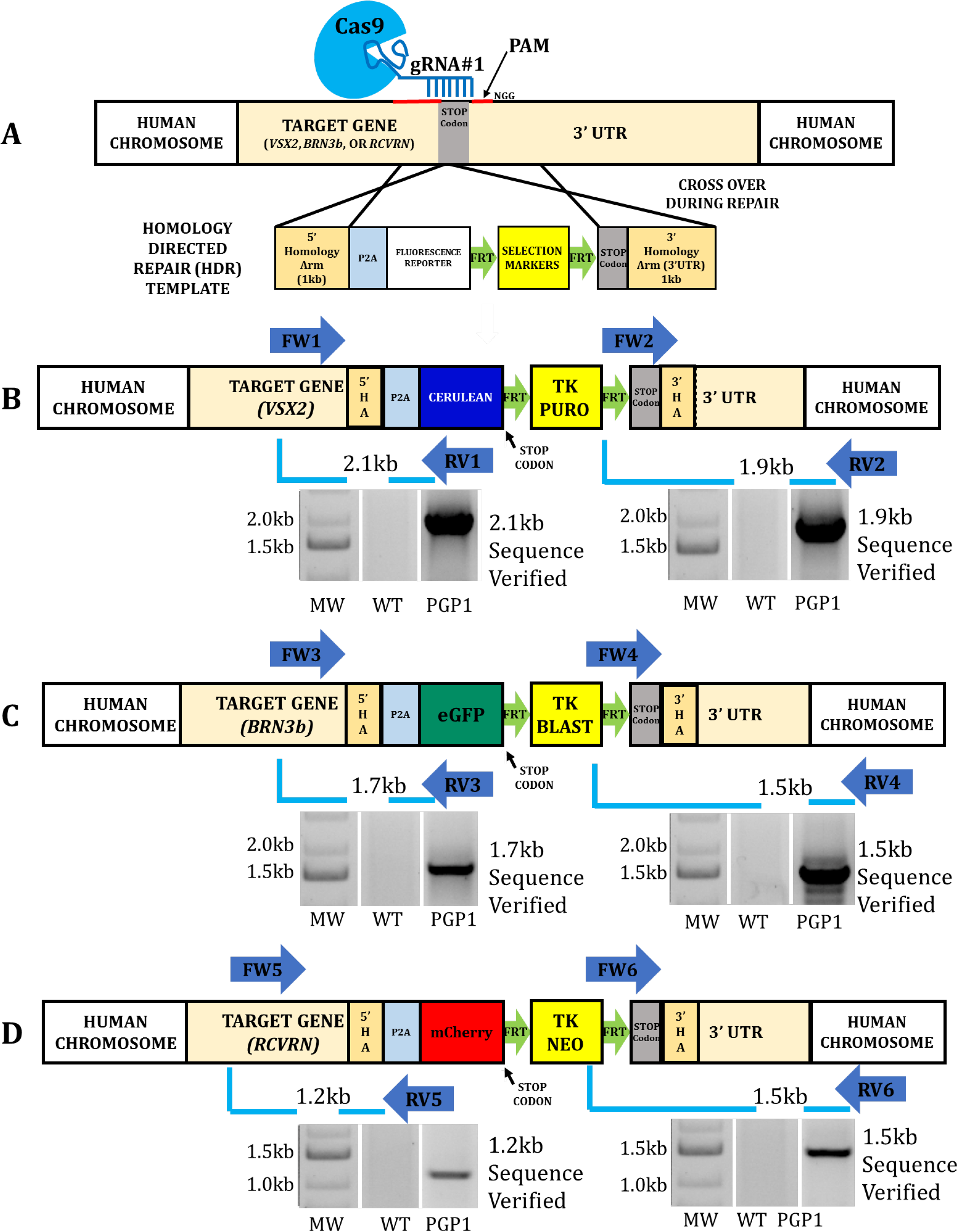
Creation of the Neural Retina Progenitor/ Retinal Ganglion Cell/ Photoreceptor (PGP1) Reporter hiPSC Line by CRISPR/Cas9 Genome Editing. (A) Schematic illustration of the generalized CRISPR/Cas9-mediated insertion strategy. CRISPR/Cas9 mediated the replacement of the endogenous STOP codon of *VSX2* (B), *BRN3b* (C), and *RCVRN* (D) loci with P2A:Cerulean, P2A:eGFP, and P2A:mCherry by homologous recombination in WT hiPSCs. Following nucleofection and triple antibiotic selection for Puromycin (PURO), Blasticidin (BLAST), and G418 (NEO) the resistant clones were screened by PCR with primer sets. (B) FW1/RV1 (forward primer (FW) located outside *VSX2* 5’HA and reverse primer (RV) located inside Cerulean) with the expected band size of 2.1kb, and FW2/RV2 (inside Puromycin to outside *VSX2* 3’HA) with the expected band size of 1.9kb. (C) FW3/RV3 (outside *BRN3b* 5’HA to inside membrane tagged enhanced GFP) with the expected band size of 1.7kb; and FW4/RV4 (inside Blasticidin to outside *BRN3b* 3’HA) with the expected band size of 1.5kb. (D) FW5/RV5 (outside *RCVRN* 5’HA to inside mCherry) with the expected band size of 1.2kb; and FW6/RV6 (inside NEO to outside *RCVRN* 3’HA) with the expected band size of 1.5kb. The WT hiPSC was used as control where no bands were seen. All the positive PCR bands were verified by sequencing. The original gels that were cropped for clarity in this figure (with white spaces between non-adjacent lanes) can be seen in their entirety in Fig. S5.

The creation of the PGP1 line involved two sequential rounds of CRISPR/Cas9 genome editing. The first step consisted of targeting the *RCVRN* locus in wild-type (WT) hiPSCs with the mCherry fluorescent protein followed by selection with G418 (Fig. 1 D). Forty-eight of the G418 resistant hiPSC clones tested by PCR analysis (blue arrows) and DNA sequencing identified three (6.25%) correctly targeted clones. Simultaneous targeting of a G418 resistant RCVRN-targeted hiPSC clone with sgRNAs and HDR targeting constructs for *VSX2* and *BRN3b* (Fig. 1 B, C) resulted in the selection (puromycin and blasticidin) of 144 resistant clones. Of these triple resistant clones, four (2.8%) were correctly targeted at the *VSX2*, *BRN3b*, and *RCVRN*, loci. Of the three triple targeted hiPSC lines isolated, we conducted our subsequent analysis on one line that we designated PGP1.

To characterize the molecular features of the PGP1 line, we used primers for PCR and DNA sequencing to analyze both alleles of the targeted genes as well as to screen for potential Cas9-generated off-target mutations. A three-primer PCR strategy, utilizing primers introduced in Fig.1, simultaneously tested for both the targeted and WT alleles. The forward primer used for screening the 5’ targeting event (Fig. 1) acted as a common primer from the endogenous gene. Two different reverse primers, one made to the fluorescent reporter gene specific for the targeted allele, and one made to the 3’ untranslated region of the endogenous gene specific for the WT allele, made it possible to determine if the clone was heterozygous or homozygous for the targeting event. PCR amplification produced DNA fragments of the expected size for both WT and targeted alleles for each gene from PGP1 genomic DNA (Fig. 2). Sequence analysis of the DNA band consistent with the WT allele from each target gene size failed to reveal any indel mutations at the sgRNA cut site (Fig. S1). DNA sequencing facilitated the analysis of PCR fragments amplified by primers surrounding predicted high-scoring off-target sites for each sgRNA. This analysis revealed no off-target mutations in PGP1 (Fig. S1 and Table S2).

**Figure 2.**
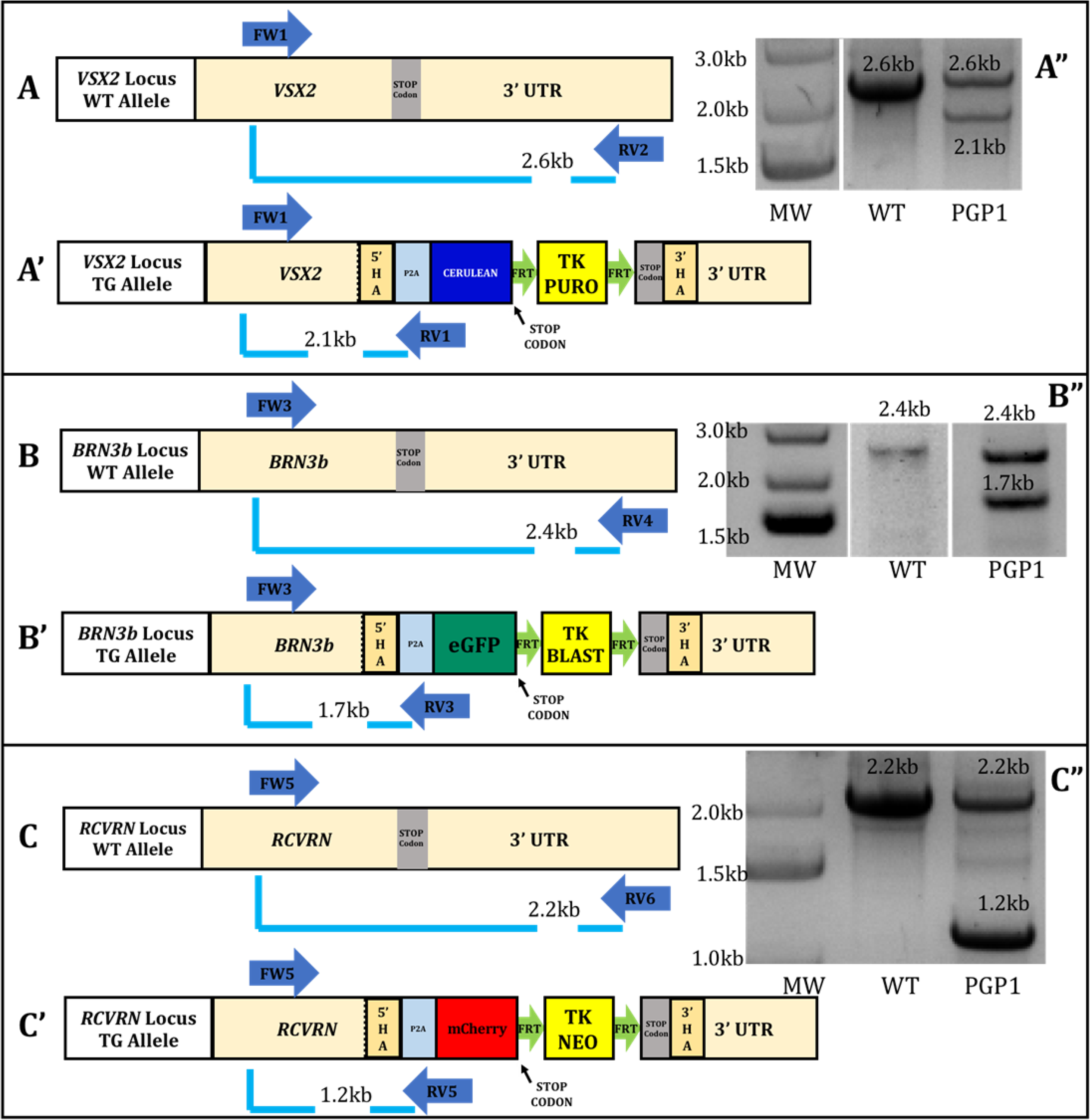
Determining the Zygosity of the PGP1 line. In each case, a three-primer PCR strategy determined whether the PGP1 clone was homozygous or heterozygous at the targeted loci. For the *VSX2* locus, primer FW1 is located outside the *VSX2* 5’HA, RV1 is located inside Cerulean, and RV2 is located outside the *VSX2* 3’HA. (A) The un-edited *VSX2* allele (FW1/RV2) should generate a 2.6kb band, (A’) while the edited *VSX2* allele (FW1/RV1) should generate a 2.1kb band. (A’’) A gel image showed bands of 2.6kb and a 2.1kb for the PGP1 line while the WT hiPSC showed a 2.6kb band indicating that PGP1 is heterozygous at the *VSX2* locus. For the *BRN3b* locus, FW3 is located outside the *BRN3b* 5’HA, RV3 is located inside the eGFP, and RV4 is located outside the *BRN3b* 3’HA. (B) The un-edited *BRN3b* allele (FW3/RV4) should generate a 2.4kb band, (B’) while the edited *BRN3b* allele (FW3/RV3) should generate a band of 1.7kb. (B”) A gel image showed bands of 2.4kb and a 1.7kb for the PGP1 line while the WT hiPSC showed only a 2.4kb band, showing that PGP1 is heterozygous at the *BRN3b* locus. For the *RCVRN* locus, FW5 is located outside the *RCVRN* 5’HA, RV5 is located inside mCherry, and RV6 is located outside the *RCVRN* 3’HA. (C) The un-edited *RCVRN* allele (FW5/RV6) should generate a 2.2kb band, (C’) while the edited *RCVRN* allele (FW5/RV5) should generate a 1.2kb band. (C”) A gel image showed bands of 2.2kb and a 1.2kb for the PGP1 line while the WT hiPSC showed only a 2.2kb band, indicating that PGP1 is heterozygous at the *RCVRN* locus. The original gels that were cropped for clarity in this figure (with white spaces between non-adjacent lanes) can be seen in their entirety in Fig. S6.

### Functional Analysis of the Fluorescent Reporters in the PGP1 Line

To confirm that all three fluorescent reporter genes would express appropriately during retinal differentiation, we created retinal organoids from the PGP1 hiPSCs. Retinal organoids were differentiated from PGP1 hiPSC-derived embryoid bodies as previously described^16^. Initially, free-floating embryoids were seeded onto Matrigel-coated plates to allow for the formation of eye field domains. At day 20 (D20) of differentiation, these eye field domains expressed Cerulean but not eGFP or mCherry (Fig. S2). After four weeks of differentiation, Cerulean positive retinal domains were manually detached from the Matrigel-coated plates and cultured as free-floating 3-dimensional retinal organoids. These organoids formed spherical cups after 55 days of differentiation (Fig. S3). By day 55 (D55) organoids revealed widespread fluorescence of Cerulean protein, expressed from the *VSX2* promoter (Fig. 3 A). Likewise, at D55 these organoids exhibited eGFP expression in a more limited population of cells, driven by the *BRN3b* promoter, and these eGFP-expressing cells preferentially localized to the interior portion of the organoids (Fig. 3 B). During construction of the BRN3b/eGFP targeting vector, we inserted a cell membrane signal peptide tag (GAP43 palmitoylation sequence) to drive membrane eGFP fluorescence^17^. This modification facilitated visualization of developing retinal ganglion cell axons at this stage. At D55, mCherry fluorescence, driven by the *RCVRN* promoter, appeared sporadically in a few cells throughout the organoids (Fig. 3 C). A merged image of all three fluorescent signals shows the restriction of fluorescent protein expression to distinct cells with little evidence of overlapping expression of different reporters (Fig. 3D). The three dimensional nature of the fluorescent protein expression in the retinal organoid can best be appreciated in a video taken from a portion of a D55 organoid (Supplemental Movie File).

**Figure 3.**
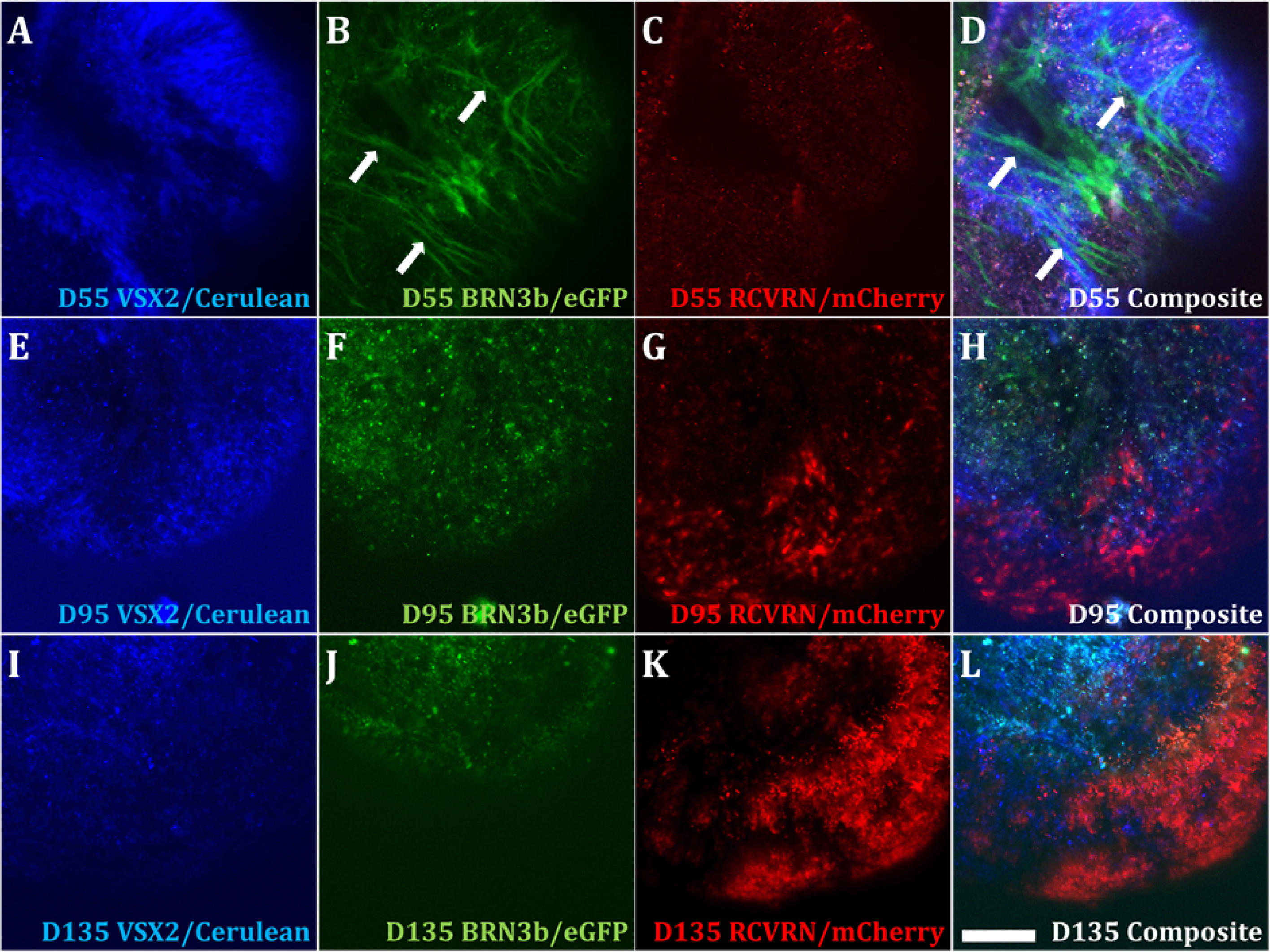
Functional Analysis of the Fluorescent Reporters in the PGP1 Line via Retinal Organoid Formation. Single three-dimensional retinal organoids derived from the PGP1 hiPSC line were visualized by fluorescent microscopy at D55 (A, B, C, D), D95 (E, F, G, H) and D135 (I, J, K, L) of differentiation. Organoids were visualized to excite Cerulean, driven by the *VSX2* promoter (A, E, I), eGFP driven by the *BRN3b* promoter (B, F J) and mCherry driven by the *RCVRN* promoter (C, G, K). Composite images represent the merger of all three fluorescent signals (D, H, L). The eGFP signal was targeted to the cell membrane by a GAP43 tag to allow for visualization of ganglion cell axons (white arrows in B, D). As organoids matured from D55 to D135, the number of cells expressing mCherry dramatically increased (compare C and K) and the eGFP positive cells populated the interior of the organoid while the mCherry positive cells occupy the organoid periphery (see J and K). Magnification Bar 100μm.

As the organoids continued to mature, layering of the fluorescent cell types became more distinct and the ratio of cells expressing different fluorescent proteins changed. At D95, Cerulean expression largely occupied an intermediate layer within the organoid (Fib. 3E, H) with eGFP concentrated on the organoid interior (Fig. 3F, H) and mCherry expressing cells organized along the organoid periphery (Fig. 3G, H). Although the number of eGFP-expressing cells increased from D55 to D95, elongated ganglion cell axons were not as distinguishable as they were at D55. By D135, the organoids had increased in size with Cerulean and eGFP positive cells occupying space in the interior of the organoid with abundant mCherry fluorescence apparent in the peripheral layers of the organoid (Fig. 3I-L).

To confirm that the observed fluorescent protein expression in PGP1-derived retinal organoids corresponded to the appropriately targeted cell types, we sorted cells from the organoids based on fluorescent protein expression. At D55, the organoids contained many cells positive for Cerulean and eGFP, but relatively few mCherry positive cells. However, by D135 the organoids contained abundant mCherry positive cells. Considering this, we performed Fluorescence Activated Cell Sorting (FACS) on proteolytically disaggregated single cell suspensions to collect Cerulean (Fig. 4 A) and eGFP (Fig. 4 D) expressing cells from D55 organoids and collected mCherry (Fig. 4 G) expressing cells from D135 organoids. Gates for collection of appropriate fluorescent cells were established by sorting human embryonic kidney (HEK293) cells transiently transfected with plasmids directing constitutive expression of Cerulean, eGFP or mCherry. At D55, Cerulean positive cells made up approximately 11.3%, and eGFP positive cells comprised approximately 4.9% of the single cells from disaggregated organoids consisting of 728,023 total cells. At D135, approximately 53% of the organoid cells (1,080,000 total cells) were mCherry positive. We also sorted organoids from wild type hiPSCs using the same gates used for the PGP1-derived organoids and found no cells within the Cerulean (Fig. 4 J), eGFP (Fig. 4 K) or mCherry (Fig. 4 L) gates.

**Figure 4.**
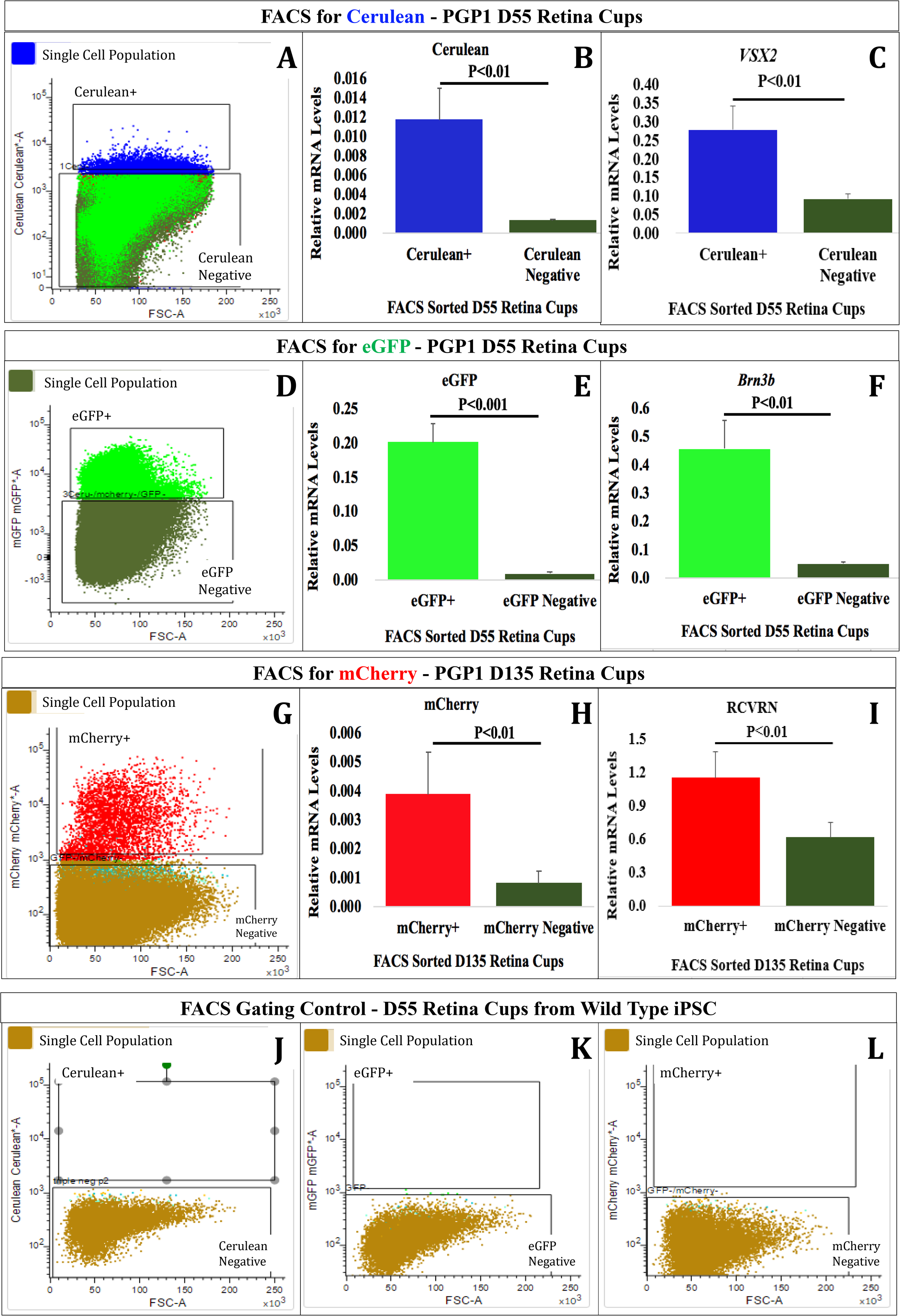
Confirmation of PGP1-derived retinal organoids corresponded to the appropriately targeted cell types. At D55 (A-F, J-L) and D135 (G-I), the retinal organoids were dissociated into single cells, and used for FACS analysis. At D55, (A) the Cerulean positive population was sorted from the Cerulean negative population. RT-qPCR analysis revealed that the Cerulean positive cells expressed significantly more Cerulean mRNA (B) and VSX2 mRNA (C) than the Cerulean negative cells. Also at D55, (D) the eGFP positive population was sorted from the GFP negative population. RT-qPCR analysis demonstrated that the eGFP positive cells expressed significantly more eGFP mRNA (E) and Brn3b mRNA (F) than the eGFP negative cells. At D135, (G) the mCherry positive population was sorted from the mCherry negative population. RT-qPCR analysis revealed that the mCherry positive cells expressed significantly more mCherry mRNA (H) and RCVRN mRNA (I) than mCherry negative cells. As a negative control, retinal organoids (D55) derived from wild-type hiPSC cells were run through the FACS and analyzed with the gates used for Cerulean (J), eGFP (K) and mCherry (L) with no cells occupying those gates. All RT-qPCR data was normalized to GAPDH expression. Error bars represent standard error of the mean (SE).

To validate our FACS gating and explore the nature of the sorted populations, we performed reverse transcription - quantitative PCR (RT-qPCR) analysis on sorted populations of cells. In each case, we sorted disaggregated organoids independently for each color into three separate tubes and compared cells positive for each sorted color to the population of cells that were negative for that sorted color according to our pre-established gates. Each of the three sorts was tested by RT-qPCR in triplicate, meaning that all graphical data represented nine RT-qPCR reactions. As expected, the Cerulean positive population expressed significantly more Cerulean mRNA than the Cerulean negative population (Fig. 4 B). Likewise, the Cerulean positive population expressed approximately 3 times more VSX2 mRNA than the Cerulean negative population (Fig. 4 C). The eGFP positive sorted population expressed 22 fold more eGFP mRNA and 9.1 fold more BRN3b mRNA than the eGFP negative population of cells (Fig. 4 E, F). On D135, the mCherry positive sorted population expressed approximately 4.9 times more mCherry mRNA and 1.9 times more RCVRN mRNA than the mCherry negative sorted population (Fig. 4 H, I). This demonstrates that the fluorescent reporters faithfully reveal the cells expressing the gene to which they were targeted. For both *VSX2* and *BRN3b* the large majority of expressing cells appeared in the Cerulean and eGFP positive populations, respectively. The appearance of relatively more RCVRN transcripts in the mCherry-negative population (although significantly less than RCVRN expression in the mCherry positive population) suggests that not all mCherry-expressing cells were captured within our gate. Alternatively, it is possible that the targeted *RCVRN* allele expresses RCVRN and mCherry less efficiently than the non-targeted *RCVRN* allele.

### PGP1-Derived Retina Organoids Contain All Major Retina Cell Types

A comparison of transcripts expressed by PGP1 hiPSCs at D0 of differentiation with those expressed by PGP1-derived retinal organoids at D55 revealed a marked decrease in pluripotency-related genes and a significant increase in retina differentiation-related genes. Pluripotency transcripts for *OCT4* (*POU5F1*) and *NANOG* expressed abundantly in PGP1-hiPSCs at D0 but virtually disappeared in the D55 organoids (Fig. 5 A). In contrast, the retinal progenitor transcripts for *PAX6*, *SIX3*, *LHX2* and *VSX2* increased significantly during early retinal organoid differentiation (Fig. 5 B). Transcripts for ganglion cells (*BRN3a* and *BRN3b*), photoreceptors (*RCVRN*) and retina pigment epithelium (*MITF* and *BEST1*) also increased significantly from D0 to D55 of differentiation (Fig. 5 C, D, E).

**Figure 5.**
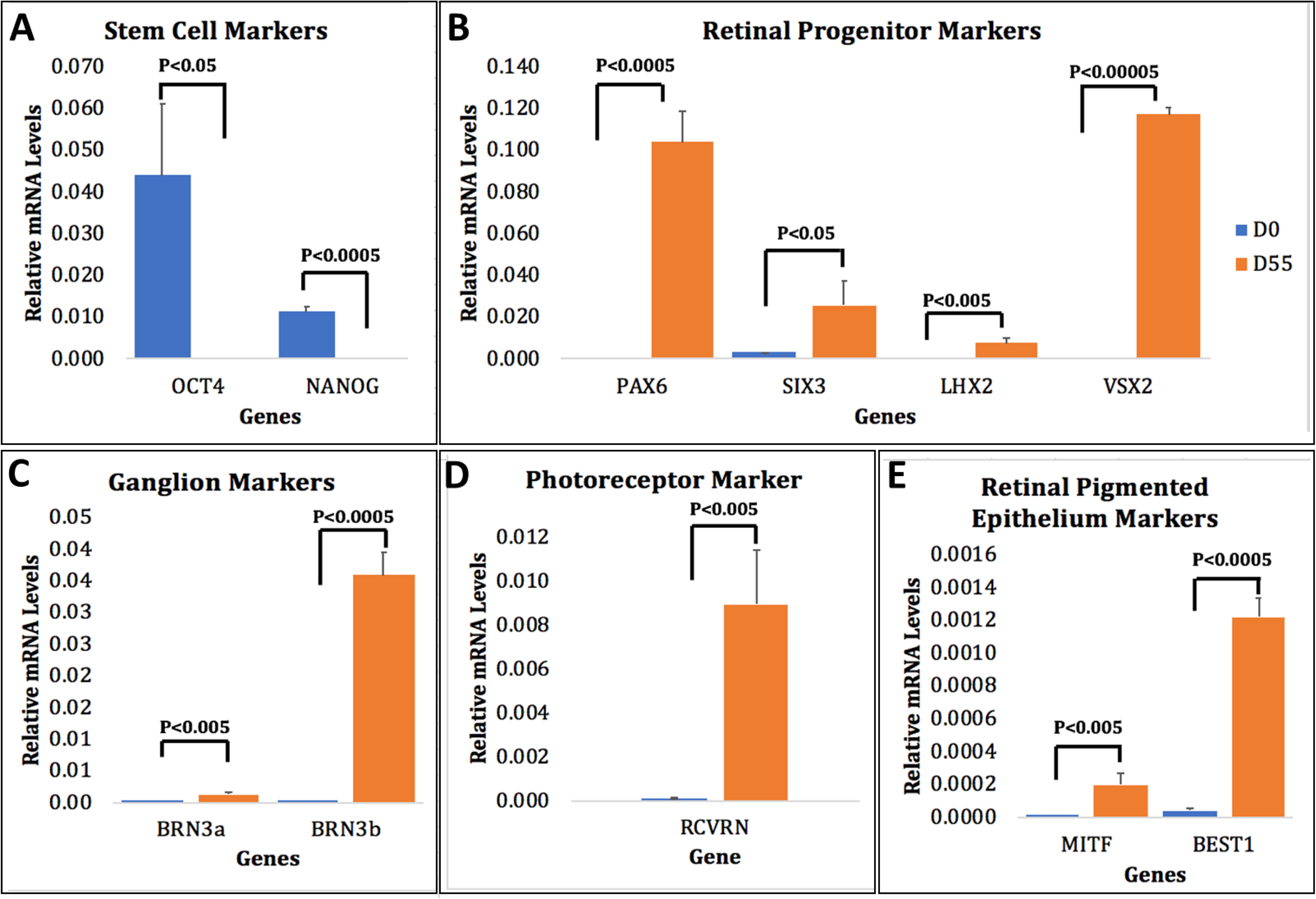
Retinal Development is Recapitulated in the Differentiating PGP1 hiPSC-Derived Retinal Organoids. Undifferentiated PGP1 hiPSCs day 0 (D0, blue) and differentiated PGP1 retinal organoids at day 55 (D55, orange) were used to measure mRNA levels of various markers via RT-qPCR normalized to GAPDH. (A) Expression of stem cell markers OCT4, NANOG; (B) retinal progenitors and neuronal markers – PAX6, SIX3, LHX2, VSX2; (C) ganglion cell markers – BRN3a, BRN3b; (D) photoreceptor marker – RCVRN; and (E) retinal pigmented epithelium markers – MITF, BEST1. As organoid differentiation progressed from D0 to D55 all stem cell markers significantly decreased and markers of retina and RPE differentiation significantly increased. Error bars represent standard error of the mean (SE).

PGP1-derived retinal organoids contained cells matching the immunological profile of all major retinal cell types. At D55, the organoids expressed the eye field precursor RX (Fig. 6 A), retinal progenitor markers: PAX6, SIX3, VSX2 (Fig. 6 B, C, D), and the proliferation marker MCM2 (Fig. 6 E). All of these aforementioned proteins appeared throughout the D55 organoid tissue. In contrast, the expression of BRN3b, a protein characteristic of retinal ganglion cells, appeared largely absent from the outer layer of the organoid, occupying a distinct localization within the organoid interior (outlined in Fig. 6 F). By D70, AP-2α, a protein expressed by amacrine cells, also appeared in the interior region of the organoid (outlined in Fig. 6 G). Although RCVRN expression, marking differentiating rods and cones, initially appeared throughout the organoid (not shown), by D70 RCVRN expressing cells appeared most prominently near the outer edge of the organoids (outlined in Fig. 6 H). At D95, PROX1 expression, characteristic of horizontal cells, occupied a space below the putative outer nuclear layer of the retina organoid (outlined in Fig. 6 I). In retinal organoids, Müller glia cells represent a late differentiating cell type. We observed CRALBP expression, characteristic of Müller glial cells, through the full thickness of the retinal organoids at D163 (Fig. 6 J). Although VSX2 expression characterizes proliferating (MCM2 positive) retina progenitor cells, VSX2 re-appears in differentiated, non-proliferating (MCM2 negative) bipolar cells by D95 (not shown). These VSX2 positive/MCM2 negative bipolar cells increase in abundance by D163 (Fig. 6 K, L).

**Figure 6.**
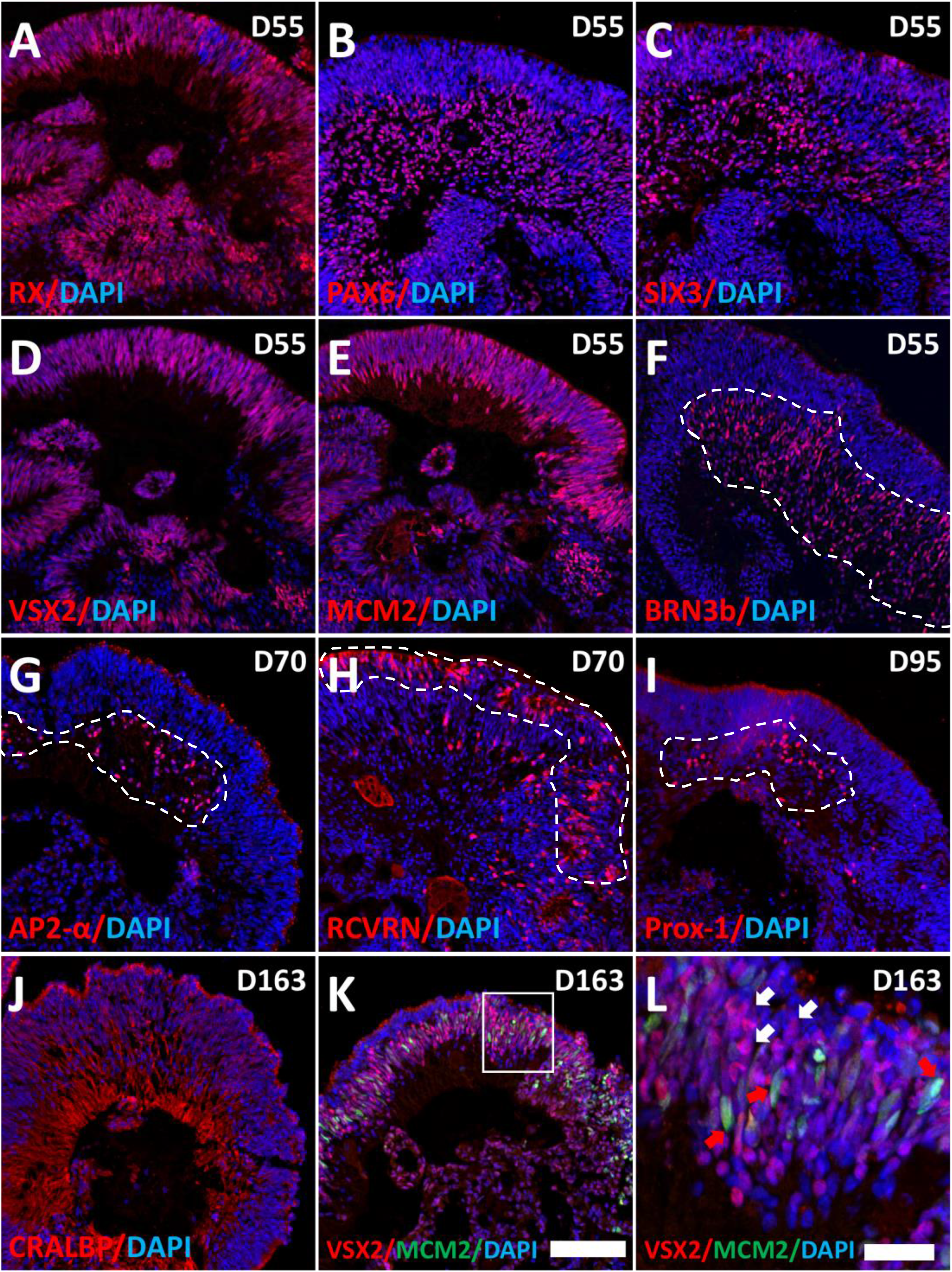
Retinal Organoids Differentiated from PGP1 Line Contain All Major Retinal Cell Types. At D55 of retinal organoid differentiation, the organoids were analyzed by immunohistochemistry using antibodies for the eye-field precursor marker RX (A), neuronal markers: PAX6 (B), SIX3 (C), the neural retina progenitor cell marker VSX2 (D), the proliferation marker MCM2 (E), and the retinal ganglion cell marker BRN3b (F). At D70 of the retinal organoid differentiation, the organoids contain the amacrine cell marker AP-2α (G) and the photoreceptor marker RCVRN (H). The organoids contain the horizontal cell marker Prox-1 (I) at D95, and the Müller glia cell marker CRALBP (J) at D163. Also at D163, the organoids contain the bipolar cell marker VSX2+/MCM2-(K, L). (L) is an enlarged view of the boxed area in (K) where VSX2+/MCM2+ (red arrows) cells represent neural retina progenitors and the VSX2+/MCM2-(white arrows) cells represent bipolar cells at D163. Dotted lines emphasize that the major expression domains of Brn3b (F), AP2α (G) and Prox1 (I) appear on the interior of the organoids while RCVRN (H) is predominantly on the organoid periphery. Magnification Bar 100μm (A-K), 25μm (L).

### DISCUSSION

Our overall goal encompassed the creation of a hiPSC line to permit real time analysis of neural retinal cell differentiation. Here, we have described a straightforward strategy to create multiple retina cell-type-specific reporters via CRISPR/Cas9 genome editing in hiPSCs. Furthermore, we functionally characterized a single, multi-targeted hiPSC clone (PGP1) both by appropriate fluorescent protein expression and by differentiation into all major retinal cell types during retinal organoid formation. Although previous publications have described single retinal cell-type-specific reporters, to our knowledge, PGP1 represents the first triple targeted retinal reporter hiPSC clone^5–7^. The ability of the PGP1 line to facilitate visual observation of retinal cell differentiation, without compromising retinal organoid formation typical of WT hiPSCs^16^, makes this clone particularly useful for a number of different applications.

The PGP1 line provides a powerful tool for real time retinal disease modeling. Retinal organoids derived from hiPSCs already play a major role in modeling eye diseases caused by known genetic causes. For example, hiPSCs derived retinal organoids from patients with mutations in genes leading to retinitis pigmentosa, and Leber congenital amaurosis have provided important insights about the pathobiology of these diseases ^18–21^. Likewise, retinal ganglion cells with specific mutations, differentiated from patient-derived hiPSCs, have provided models for optic atrophy and glaucoma^22,23^. The relative ease of creating specific mutations with CRISPR/Cas9 genome editing should make it possible to take advantage of the cell-type-specific reporters in PGP1 when modeling virtually any genetic retinal disease.^19^

PGP1-derived retinal organoids could also provide a platform to test drugs or chemicals to treat various retina diseases or for retinal toxicity. A number of different physical or chemical insults can result in photoreceptor damage, making the search for compounds that can prevent or minimize such damage of great importance for preserving vision. Retinal organoids made from mouse iPSCs, engineered with an Nrl-eGFP reporter to label rod photoreceptors, facilitated the testing of compounds to protect photoreceptors from 4-hydroxytamoxifen-induced degeneration^24^. The PGP1 hiPSC line could easily be used in a similar way with the added advantage of simultaneously monitoring retinal progenitors, bipolar cells and ganglion cells. The recently reported three-dimensional automated reporter quantification (3D-ARQ) technology^25^ should make screening drugs that affect retina development and/or survival straightforward in PGP1-derived retinal organoids. In many cases, the development of drugs to treat specific diseases of the retina suffer from the inability to purify large numbers of specific retina cell types. For example, drugs to treat AMD or glaucoma could benefit from methods that achieve pure populations of photoreceptors or ganglion cells, respectively.

The inherent reporters in the PGP1 line will facilitate optimization of protocols to achieve the differentiation and/or survival of specific retinal cell types. Despite the advances that led to retinal organoid formation from hiPSCs, there clearly exists room for improvement in organoid technology. Improved mature photoreceptor yields and laminar stratification resulted from the use of stirred-tank bioreactors, most likely due to increased cell proliferation and decreased apoptosis^25^. This particular study relied on immunohistochemical characterization and FACS-mediated immuno purification of organoid-derived cell types. However, PGP1-derived retinal organoids would permit similar protocol optimization for multiple cell types in real time without relying on antibody labels. The ability to monitor particular cell types continuously without interruption should improve optimization for the survival of desired cells. Specifically, one of the greatest challenges with current retinal organoid technology is achieving long-term survival of retinal ganglion cells.

The PGP1 line could provide an effective strategy for improved purification of specific cell types, either from retinal organoids or from protocols designed to achieve direct retinal cell type differentiation. During the creation of hiPSC-derived retinal organoids and during normal retinal development, retinal cell types develop precisely in concert. Even in direct differentiation protocols to achieve a particular retinal cell type from hiPSCs, purifying fully differentiated, mature cell types from partially differentiated progenitors remains an issue^26^. The purification of specific retinal neuron cell types can often facilitate downstream analysis, drug screening, or isolation of particular cell populations for transplantation studies. Recombinant AAV viruses that contained a GFP reporter driven by an L/M cone-specific pR2.1 promoter led to the successful isolation of cone photoreceptors following FACS from both human fetal retina and from hiPSC-derived retinal organoids^27^. A similar strategy used recombinant lentivirus to achieve GFP expression from the photoreceptor-specific (IRBP) promoter to achieve FACS purification of hiPSC-derived photoreceptor cells prior to transplantation into immunocompromised mice^28^. Although both of these reports relied on virus-mediated reporter gene expression, others have adopted the use of endogenous reporters. For example, hiPSCs engineered with a P2A peptide-mCherry reporter into the *BRN3b* locus made it possible to purify viable retinal ganglion cells from retinal organoids by FACS^5^. These examples all attest to the usefulness of the PGP1 line for purification of specific retinal cell types. However, the unique feature of the PGP1 line versus previously reported lines lies in its potential for simultaneously purifying retinal progenitor cells, retinal ganglion cells, photoreceptors and bipolar cells.

While fluorescently labeled cells would presumably not be used directly in therapies, a clone with multiple cell-type-specific reporters could serve as a powerful tool for proof-of-concept studies and for tracking retinal specific cell fate post-transplantation in animal models. Cell-based transplantation approaches for the treatment of glaucoma and other ocular neurodegenerative disorders already exist. For example, transplanting human neural progenitor cells into a mouse glaucoma model increased host retinal ganglion cell survival^29^. Although the transplanted cells restored some vision, the mechanism that led to vision improvement by these cells remains unclear. In the future, PGP1-derived cells will make it possible to evaluate the fate of transplanted cells *in vivo*. In these transplants, PGP1 progenitors, and later bipolar cells, would exhibit blue fluorescence, while ganglion cells or photoreceptors derived from the transplant would exhibit green or red fluorescence, respectively. In another study, transplantation of GFP-labeled human photoreceptors into a rat model of retinitis pigmentosa demonstrated restoration of the host rod function^30^. Although these authors reported integration of GFP positive photoreceptors into the outer nuclear layer, the recent realization of cytoplasmic transfer from donor to host cells in photoreceptor transplantation^31,32^ necessitates further confirmation of donor cell integration. mCherry positive photoreceptors from the PGP1 line could provide utility to not only repeat these experiments, but to also serve as an indicator of both cell survival and cell fate of the transplanted photoreceptors.

In summary, we demonstrate a CRISPR/Cas9 strategy to target the expression of multiple fluorescent reporter genes into endogenous loci in hiPSCs. In doing so, we created a triple transgenic hiPSC line (PGP1) and tested the function of this line by directed differentiation into 3D retinal organoids. Organoids produced from the PGP1 line expressed Cerulean in neural retina progenitors and bipolar cells, membrane-targeted eGFP in retinal ganglion cells and mCherry in photoreceptors. In addition, PGP1-derived retinal organoids contained all major retinal cell types. The usefulness of this strategy extends to virtually any cell-type-specific gene, limited only by the number of different fluorescent reporters available for simultaneous analysis. The PGP1 line, and subsequent hiPSC lines developed using this approach, hold great promise for studying retinal development, disease modeling, drug screening, and pre-clinical transplantation studies.

## METHODS

### hiPSC Culture

Human umbilical cord stem cell derived hiPSC6.2^33^ (Life Technologies, A18945) were grown on Matrigel-coated (MG) plates using chemically defined Essential 8 medium (Thermo fisher, A1517001) as described previously^34^. The medium was changed daily, and cells were passaged every 3-4 days using 0.5mM EDTA in 1xDPBS without calcium and magnesium to lift cells from the tissue culture plate (Thermo fisher, 15575020).

### Specific Single Guide RNA Vector (sgRNA) Design

CRISPR specific guide RNAs (sgRNAs) (Table S3) were individually cloned into a U6-driven sgRNA expression PX458 vector that includes the *S. pyogenes* Cas9 coding sequence (Addgene, #48138) as described^35^. To determine the Cas9 cutting efficiency of these sgRNAs, we transfected each vector into Human Embryonic Kidney (HEK293) cells. Forty-eight hours after transfection, HEK293 DNA was extracted and amplified by Q5 High-Fidelity PCR (NEB, M0494S) using primers encompassing the sgRNA recognition site. The PCR products were digested with T7 Endonuclease I (T7E1, NEB M0302S) according to the recommended protocol^36^. Successful Cas9 cleavage by sgRNAs resulted in two distinct bands in the T7E1 assay.

### Homology Directed Repair (HDR) Template Generation

To generate the HDR templates, the left and right homology arms (HA) of each locus were amplified from the WT hiPSC genomic DNA using primers listed in Table S4. The amplified homology arms were inserted into the P2A:Cerulean.pL451; P2A:GAP43.eGFP.pL451, P2A:mCherry.pL451 vectors via Gibson Assembly (NEB^37^). The Cerulean tag came from addgene #53749, the membrane bound form of eGFP from addgene #4757, and mCherry from addgene #26901. DNA sequencing verified the generated HDR templates following PCR amplification.

### Insertion of Fluorescent Reporter Genes into Selected Loci

To generate the RCVRN/mCherry hiPSC lines we transfected 2.5μg of the HDR template and 2.5μg of the sgRNA vector into WT hiPSCs using a 4D-Nucleofector X Unit and the P3 Primary Cell kit (Amaxa, V4XP-3012) based on manufacturer’s protocol. After transfection, the cells were cultured with Essential 8 media plus 10μM ROCK inhibitor (Sigma SCM075) overnight. Subsequently, culture media was changed daily without addition of ROCK inhibitor. Forty-eight hours after transfection, antibiotic selection began with 100 μg/ml and slowly increased to 250 μg/ml of G418 (Corning, 30234CR) over the course of one week to select for Neomycin resistant colonies. Resistant clones were manually picked and cultured individually in a 48 well plate. The DNA of each clone was extracted (Zymo Research, D3025), and screened for reporter integration by PCR (Thermo Fisher, EP1701) using the primers listed in Table S4. The clones, which exhibited the expected PCR fragment sizes on each side of the HDR junctions, were validated by DNA sequencing. To generate the triple transgenic hiPSCs, a RCVRN/mCherry hiPSC clone was transfected with 1.25ug each of: the *BRN3b/*eGFP HDR template, the sgRNA vector to the *BRN3b* locus, the *VSX2*/Cerulean HDR template, and the sgRNA vector to the *VSX2* locus using nucleofection as described above. Double antibiotic selection was performed using 100 μg/ml Blasticidin and 100 μg/ml Puromycin, and slowly increased up to 250 μg/ml of each antibiotic over the course of one week to select for Blasticidin and Puromycin resistant colonies. Resistant clones were manually picked and cultured in a 48 well plate (one clone per well). Again, DNA from each clone was extracted and screened by PCR using primers listed in Table S4. PCR bands of the expected size were verified by DNA sequencing.

### Off-Target Screening

The PGP1 clone was screened by selecting five high scoring off-target sites for each sgRNA used according to online tools provided by Benchling^38^. Each potential sgRNA off target site listed in Table S2 (off-target score provided by Benchling) was screened by High-Fidelity PCR (Q5 NEB, M0491L) with primers listed in Table S5 and PCR products were sequenced using Eurofins Genomics tube DNA sequencing services. Each result was independently repeated five times.

### 3-D Retinal Organoid Generation from the PGP1 Triple Targeted Line

The PGP1 line was used to create 3-D retinal organoids as described previously using the Zhong et al., 2014 protocol with the following modifications^16^. Briefly, hiPSC were incubated in 0.5 mM EDTA/DPBS (Thermo fisher, 15575020) for 5 minutes at 37° C. Cells were then dissociated into small clumps and cultured in mTeSR1 medium with 10 μM ROCK inhibitor to form aggregates. The aggregates were gradually transitioned into neural induction medium (NIM)^16^ for three days (D1-3 of differentiation), then cultured in NIM from D3 to D6. On D7, the aggregates were seeded on Matrigel (hESC-qualified, Corning)-coated dishes in NIM at an approximate density of 20 aggregates per cm^2^ and switched to DMEM/F12 (3:1) supplemented with 2% Gem21 NeuroPlex (without vitamin A, Gemini Bio-Products, 400-161), 1X NEAA, and 1% antibiotic-antimycotic (Thermo, 15240062) on D16. The medium was changed every 3 days. On the fourth week (D28) of differentiation, a cell scraper was used to detach the cells from the dishes and and the cells were transferred to petri dishes. The cells were then cultured in suspension at 37°C in a humidified 5% CO_2_ incubator in DMEM/F12 (3:1) supplemented with 2% Gem21 NeuroPlex, 1X NEAA, and 1% antibiotic-antimycotic. Within 3-5 days, cells began forming 3-D retinal organoids. The organoids were then mechanically separated from the rest of the cells using sharpened tungsten needles under a dissecting microscope. From that point on, the medium was changed twice a week. To culture the retinal organoids long-term, the medium was supplemented with 10% fetal bovine serum (Gibco), and 2 mM GlutaMax (Invitrogen) beginning on D42.

### Reverse Transcription-PCR Analysis of Retinal Organoids

Total RNA isolation of hiPSC colonies or retinal organoids was done using either Quick RNA miniprep plus (Zymo research, R1057) or 96-well plate RNA extraction kit (Illustra RNAspin 96, 25-0500-75) using the manufacturer’s protocol. Reverse transcription was performed using the ImProm-II Reverse Transcription System (Promega, A3800) according to the manufacturer’s protocol. RT-qPCR was performed with GoTaq® qPCR Master Mix for Dye-Based Detection (Promega, A6001) using a CFX Connect Bio-Rad qPCR System. Forty cycles were run at 95 °C denaturation for 40 s, at 60 °C annealing for 40 s and at 72 °C extension for 60 s, using primers listed in Table S6. The expression levels of individual genes were normalized to GAPDH mRNA levels and analyzed using the detla-delta Ct method (Applied Biosystems) with significant differences revealed by a two tailed Student’s t-test. Error bars in each figure represents the standard error (SE) of three individual experiments.

### Immunofluorescence of the Retinal Organoids

Retinal organoids were fixed with 4% paraformaldehyde (4 % PFA) for 20 minutes at room temperature. The fixed organoids were incubated in 30% sucrose overnight at 4°C before embedding in OCT. Organoids were cryosectioned at 15 μm prior to immunofluorescence. Immunofluorescence was performed using antibodies specifically against the proteins of interest listed in Table S7. Fluorescent images were acquired with a Nikon Eclipse 80i microscope and/or Zeiss LSM 710 Laser Scanning Confocal System. Fluorescent protein fluorescence does not survive our fixation and embedding protocol and therefore provides no interference with secondary antibody fluorescence (Fig. S7, S8).

### Fluorescence-Activated Cell Sorting (FACS) Analysis of the Retinal Organoids

Thirty organoids each from D55 and D135 were incubated at 37° C in Accutase (Innovative Cell Technologies, AT104) for 20 minutes, broken down to single cells by pipetting, and filtered through a 40μm strainer (Fishersci, 22-363-547) to eliminate cell clumps, and resuspended in ice cold 5%FBS/1xHBSS (Fishersci, 14-025-092) at a concentration of 10 million cells/ml. These cells were filtered again through a second strainer (Fisher, 08-771-23) prior to sort. For the D55 sort, single cells were sorted to separate the Cerulean positive from the Cerulean negative population, and the eGFP positive from the eGFP negative population using a FACSMelody Cell Sorter (BD Biosciences). For the D135 sort, single cells were sorted to separate the mCherry positive from the mCherry negative population. The sorted populations were used to measure mRNA transcripts for *Cerulean*, *VSX2*, *mCherry*, *RCVRN*, *eGFP*, *BRN3b*, and *GAPDH* via RT-qPCR using the primers listed in Table S8. Gates for sorting disaggregated retinal organoids were established in a previous experiment using HEK293 cells transiently transfected with and expression plasmid for Cerulean, eGFP, or mCherry. FACS was conducted 48 hours after transfection to determine the appropriate gates for each fluorescent protein (Fig. S4). In addition, organoids produced from wild-type hiPSCs were sorted using these established gates as a negative control.

## Supporting information

3D organoid movie

## AUTHOR CONTRIBUTION

P.T.L. designed and conducted all CRISPR/Cas9 targeting and screening of hiPSC clones, performed all immunohistochemistry and confocal microscopy, designed and conducted all RT-qPCR analysis and FACS sorting. C.G. conducted all retinal organoid cultures and assisted with immunohistochemistry. K.D.R.T. participated in experimental design and reviewed all of the data for the manuscript. M.L.R. supervised all experiments, participated in all aspects of experimental design, and reviewed all the data. P.T.L. and M.L.R. wrote the manuscript. All authors reviewed and edited the manuscript.

## ACKNOWLEDGMENTS

This work was supported by grants from the Sigma Xi- Scientific Research Society (P.T.L. G201510151661206, G2017031589218117, G2018031589218117, Miami University Academic Challenge (P.T.L), Miami University College of Arts and Science (K.D.R.T. and M.L.R.), The James and Beth Lewis Endowed Professorship (M.L.R.), and National Eye Institute grant EY026816 (K.D.R.T). The FACS-Melody system was acquired with an NSF MRI grant 1726645 (M.L.R.). We would like to acknowledge Brad D. Wagner for technical assistance, and Dr. Timothy Wilson for assistance with FACS analysis. The authors acknowledge the assistance of both the Center for Bioinformatics and Functional Genomics and the Center for Advanced Microscopy and Imaging at Miami University. We would like to thank Stephanie Padula, Dr. Anthony Sallese, Nathan Burns and Adam LeFever for editing the manuscript prior to submission.

## ADDITIONAL INFORMATION

## DATA AVAILABILITY STATEMENT

The authors will make all relevant data not included with the manuscript and supplemental materials freely available upon request.

## REFERENCES

1. Sowden, J. C. ESC-Derived Retinal Pigmented Epithelial Cell Transplants in Patients: So Far, So Good. Cell Stem Cell 15, 537–538 (2014).

2. Veleri, S. et al. Biology and therapy of inherited retinal degenerative disease: insights from mouse models. Dis. Model. Mech. 8, 109–129 (2015).

3. Llonch, S., Carido, M. & Ader, M. Organoid technology for retinal repair. Dev. Biol. 433, 132–143 (2018).

4. DiStefano, T. et al. Accelerated and Improved Differentiation of Retinal Organoids from Pluripotent Stem Cells in Rotating-Wall Vessel Bioreactors. Stem Cell Reports 10, 300–313 (2018).

5. Sluch, V. M. et al. Differentiation of human ESCs to retinal ganglion cells using a CRISPR engineered reporter cell line. Sci. Rep. 5, 16595(2015).

6. Sluch, V. M. et al. Enhanced Stem Cell Differentiation and Immunopurification of Genome Engineered Human Retinal Ganglion Cells. Stem Cells Transl. Med. 6, 1972–1986 (2017).

7. Phillips, M. J. et al. Generation of a rod-specific NRL reporter line in human pluripotent stem cells. Sci. Rep. 8, (2018).

8. Shirai, H. et al. Transplantation of human embryonic stem cell-derived retinal tissue in two primate models of retinal degeneration. Proc. Natl. Acad. Sci. U. S. A. 113, E81–90 (2016).

9. Jin, Z.-B. et al. Modeling retinal degeneration using patient-specific induced pluripotent stem cells. PLoS One 6, e17084 (2011).

10. Zou, C. & Levine, E. M. Vsx2 controls eye organogenesis and retinal progenitor identity via homeodomain and non-homeodomain residues required for high affinity DNA binding. PLoS Genet. 8, e1002924 (2012).

11. Phillips, M. J. et al. Modeling human retinal development with patient-specific induced pluripotent stem cells reveals multiple roles for visual system homeobox 2. Stem Cells 32, 1480–1492 (2014).

12. Badea, T. C., Cahill, H., Ecker, J., Hattar, S. & Nathans, J. Distinct roles of transcription factors brn3a and brn3b in controlling the development, morphology, and function of retinal ganglion cells. Neuron 61, 852–864 (2009).

13. Polans, A. S. et al. Recoverin, a photoreceptor-specific calcium-binding protein, is expressed by the tumor of a patient with cancer-associated retinopathy. Proc. Natl. Acad. Sci. U. S. A. 92, 9176–9180 (1995).

14. Cao, D. & Barrionuevo, P. A. Estimating photoreceptor excitations from spectral outputs of a personal light exposure measurement device. Chronobiol. Int. 32, 270–280 (2015).

15. Wang, Y., Wang, F., Wang, R., Zhao, P. & Xia, Q. 2A self-cleaving peptide-based multi-gene expression system in the silkworm Bombyx mori. Sci. Rep. 5, 16273(2015).

16. Zhong, X. et al. Generation of three-dimensional retinal tissue with functional photoreceptors from human iPSCs. Nat. Commun. 5, 4047(2014).

17. Gauthier-Kemper, A. et al. Interplay between phosphorylation and palmitoylation mediates plasma membrane targeting and sorting of GAP43. Mol. Biol. Cell 25, 3284–3299 (2014).

18. Parfitt, D. A. et al. Identification and Correction of Mechanisms Underlying Inherited Blindness in Human iPSC-Derived Optic Cups. Cell Stem Cell 18, 769–781 (2016).

19. Schwarz, N. et al. Arl3 and RP2 regulate the trafficking of ciliary tip kinesins. Hum. Mol. Genet. 26, 2480–2492 (2017).

20. Tucker, B. A. et al. Patient-specific iPSC-derived photoreceptor precursor cells as a means to investigate retinitis pigmentosa. Elife 2, e00824 (2013).

21. Megaw, R. et al. Gelsolin dysfunction causes photoreceptor loss in induced pluripotent cell and animal retinitis pigmentosa models. Nat. Commun. 8, 271(2017).

22. Chen, J., Riazifar, H., Guan, M.-X. & Huang, T. Modeling autosomal dominant optic atrophy using induced pluripotent stem cells and identifying potential therapeutic targets. Stem Cell Res. Ther. 7, 2(2016).

23. Teotia, P. et al. Modeling Glaucoma: Retinal Ganglion Cells Generated from Induced Pluripotent Stem Cells of Patients with SIX6 Risk Allele Show Developmental Abnormalities. Stem Cells 35, 2239–2252 (2017).

24. Ito, S.-I., Onishi, A. & Takahashi, M. Chemically-induced photoreceptor degeneration and protection in mouse iPSC-derived three-dimensional retinal organoids. Stem Cell Res. 24, 94–101 (2017).

25. Ovando-Roche, P. et al. Use of bioreactors for culturing human retinal organoids improves photoreceptor yields. Stem Cell Res. Ther. 9, 156(2018).

26. Tano, K. et al. A novel in vitro method for detecting undifferentiated human pluripotent stem cells as impurities in cell therapy products using a highly efficient culture system. PLoS One 9, e110496 (2014).

27. Welby, E. et al. Isolation and Comparative Transcriptome Analysis of Human Fetal and iPSC-Derived Cone Photoreceptor Cells. Stem Cell Reports 9, 1898–1915 (2017).

28. Lamba, D. A. et al. Generation, purification and transplantation of photoreceptors derived from human induced pluripotent stem cells. PLoS One 5, e8763 (2010).

29. Ma, J. et al. Transplantation of Human Neural Progenitor Cells Expressing IGF-1 Enhances Retinal Ganglion Cell Survival. PLoS One 10, e0125695 (2015).

30. Jayaram, H. et al. Transplantation of photoreceptors derived from human Muller glia restore rod function in the P23H rat. Stem Cells Transl. Med. 3, 323–333 (2014).

31. Nickerson, P. E. B., Ortin-Martinez, A. & Wallace, V. A. Material Exchange in Photoreceptor Transplantation: Updating Our Understanding of Donor/Host Communication and the Future of Cell Engraftment Science. Front. Neural Circuits 12, 17(2018).

32. Ortin-Martinez, A. et al. A Reinterpretation of Cell Transplantation: GFP Transfer From Donor to Host Photoreceptors. Stem Cells 35, 932–939 (2017).

33. Burridge, P. W. et al. A universal system for highly efficient cardiac differentiation of human induced pluripotent stem cells that eliminates interline variability. PLoS One 6, e18293 (2011).

34. Chen, G. et al. Chemically defined conditions for human iPSC derivation and culture. Nat. Methods 8, 424–429 (2011).

35. Ran, F. A. et al. Genome engineering using the CRISPR-Cas9 system. Nat. Protoc. 8, 2281–2308 (2013).

36. Vouillot, L., Thélie, A. & Pollet, N. Comparison of T7E1 and Surveyor Mismatch Cleavage Assays to Detect Mutations Triggered by Engineered Nucleases. G3: Genes|Genomes|Genetics 5, 407–415 (2015).

37. Liu, P., Jenkins, N. A. & Copeland, N. G. A highly efficient recombineering-based method for generating conditional knockout mutations. Genome Res. 13, 476–484 (2003).

38. Benchling · Better tools, faster research. Available at: https://benchling.com/crispr. (Accessed: 8th February 2018).

