## supplemental files for "Generation of a Retina Reporter hiPSC Line to Label Progenitor, Ganglion, and Photoreceptor Cell Types"

Supplemental Table S1: Primers used for genotyping in figures 1 and 2

| **Location** | **Forward (5'-3')** | **Reverse (5'-3')** | **Size** |
|---|---|---|---|
| Outside *VSX2* 5'HA to Cerulean | CCAAGTGGAGGAAGCGGGAGAAGT (FW1) | CGGCGGCGGTCACGAAC (RV1) | 2053bp |
| Puro to outside *VSX2* 3'HA | GCGTTGGCTACCCGTGAT (FW2) | GCCCCAGCTCCTTATTCC (RV2) | 1870bp |
| Outside *BRN3b* 5'HA to eGFP | TATTCGGCGGGCTGGATGAGAGTC (FW3) | GCCGTCGCCGATGGGGGTGTT (RV3) | 1673bp |
| Blas to outside *BRN3b* 3'HA | TCGACTAGAGCTTGCGGAACC (FW4) | AACCAGGCCATATACAGAACTCAA (RV4) | 1528bp |
| Outside *RCVRN* 5'HA to mCherry | AGCTTTGTTGAGCACCGACT(FW5) | GTTCTCCTCCACGTCTCCAG (RV5) | 1167bp |
| Neo to outside *RCVRN* 3'HA | TCGCCTTCTTGACGAGTTCT (FW6) | TGGATCTGGTCCTCTCCATC (RV6) | 1493bp |
| Outisde *VSX2* 5'HA to outside *VSX2* 3'HA | CCAAGTGGAGGAAGCGGGAGAAGT (FW1) | GCCCCAGCTCCTTATTCC (RV2) | 2627bp |
| Outisde *BRN3b* 5'HA to outside *BRN3b* 3'HA | TATTCGGCGGGCTGGATGAGAGTC (FW3) | AACCAGGCCATATACAGAACTCAA (RV4) | 2438bp |
| Outisde RCVRN 5'HA to outside *RCVRN* 3'HA | AGCTTTGTTGAGCACCGACT(FW5) | TGGATCTGGTCCTCTCCATC (RV6) | 2221bp |

Supplemental Table S2: Primers used for CRISPR off-target screening

| **Off-Target Screening VSX2-sgRNA** | | | | |
|---|---|---|---|---|
| **Name** | **Gene** | **Sequence** | **PAM** | **Off-target Score*** |
| VSX2, Chr14 | | GTCAAGGCGCGCTCAGATGC | CGG | 100 |
| Chr19 non-gene sequence | | GTCAAGGCGTACTCAGATGC | GAG | 2.668116758 |
| Chr19 non-gene sequence | | GTGAAGAAGTGCTCAGATGC | CAG | 0.916843223 |
| Sytabulin, Chr8 | ENSG00000147642 | GTGAAGACACCCTCAGATGC | TGG | 0.349165048 |
| VEGF-A, Chr6 | ENSG00000112715 | GTCAAGGCGTGCTCCGATGG | GGG | 0.317986706 |
| KIF16B, Chr20 | ENSG00000089177 | GTCGAAGCGGGCTCCGATGC | AGG | 0.252128661 |
| STAT2, Chr12 | ENSG00000170581 | GTCAATGGGAGCTCTGATGC | AGG | 0.234357477 |

| **Off-Target Screening BRN3b-sgRNA** | | | | |
|---|---|---|---|---|
| **Name** | **Gene** | **Sequence** | **PAM** | **Off-target Score*** |
| BRN3b (POU4F2), Chr4 | ENSG00000151615 | AAGAGTCTTCTAAATGCCGG | CGG | 100 |
| RP11-1100L3.7, Chr12 | ENSG00000257663 | AGCAGTCTTCCAGATGCCGG | CAG | 0.371654759 |
| RP11-45M11.7, Ch6 | ENSG00000275846 | AAGCTCCTTCTAAATGCCAG | TAG | 0.351773802 |
| TC2N, Chr14 | ENSG00000165929 | TAAAGTCTTCTAAATGCCAA | TAG | 0.331943062 |
| FAM83F, Chr22 | ENSG00000133477 | AAGAGAATTGGAAATGCCGG | CAG | 0.302192873 |
| RP11-484K9.4, Chr3 | ENSG00000272844 | AAGACTCTTTGAAATGCCTG | CGG | 0.288247111 |
| RP11-321M21.1 , Chr18 | ENSG00000266774 | AATAGTCCTCCAAATGCTGG | CAG | 0.202171083 |

| **Off-Target Screening RCVRN-sgRNA** | | | | |
|---|---|---|---|---|
| **Name** | **Gene** | **Sequence** | **PAM** | **Off-target Score*** |
| Recoverin, Chr17 | ENSG00000109047 | AGGGAGGACAGCTGAACAGT | TGG | 100 |
| Chr4 non-gene sequence | | AGGGAGGCCAGCTGAAGAGT | GGG | 3.099576271 |
| Chr2 non-gene sequence | | GGAGAGGGCAGCTGAACAGT | TAG | 2.726928675 |
| Chr14 non-gene sequence | | AGAGAGATCAGCTGAACAGT | GGG | 1.740860136 |
| Chr17 non-gene sequence | | AGAAAGGACAGCTGAACTGT | AGG | 0.741790707 |
| Chr17 non-gene sequence | | AGTGAGGATAGCTGGACAGT | AGG | 0.541032634 |

* Off-target scores provided by Benchling.

Supplemental Table S3: sg RNA sequences for targeting

| **Specific guide RNA** | **5' - 3'** |
| --- | --- |
| *VSX2* .sgRNA | GTCAAGGCGCGCTCAGATGC |
| *BRN3b* .sgRNA | AAGAGTCTTCTAAATGCCGG |
| *RCVRN* .sgRNA | AGGGAGGACAGCTGAACAGT |

Supplemental Table S4: Primers used for Gibson Assembly to make the HDR template

| Gene | Forward (5’-3’) | Reverse (5’-3’) |
|---|---|---|
| VSX2.5’HA | ATTGGGTACCGGGCCTCCTGTGAGAACAGTGTG | CCGCTTCCGTCGACCAAAGCCATGTCCTCCAGC |
| VSX2.3’HA | ATACGAAGTTATTAGGTGTAGGTCAAGGCGCGCTCA | CTCCACCGCGGTGGCGCCAGATTGGGTTGTTCAAGG |
| RCVRN.5’HA | CTATAGGGCGAATTGGGTACTGCCTTCCCCGCCAGGTC | GTCGACCTCGAGGGGGGGCCTGGCGTTCTTCATCTTTTCCTTCACTTTTTG |
| RCVRN.3’HA | ATACGAAGTTATTAGGTGTGAACACACATGCACACA | CTCCACCGCGGTGGCCAAAAGCTTATTCATCGGG |
| BRN3b.5’HA | GGCGAATTGGAGCTCCACCGCGGTGGCCGCCGAGGCTCTGGCAGC | ATACAGCACAGCATAGGTCCAGGGTTCTCCTCCACG |
| BRN3b.3’HA | CCACTAGTTCTAGAAATAGAAGACTCTTGGCCTCTCC | TTGATATCGAATTCCTGCAGCCCGGGGTGCATCGGTCATGCTTCC |

Supplemental Table S5: Primers flanking the sgRNA cut site used to generate PCR fragments for sequencing

| Set of primers used to screen for off-target cutting efficiency of **VSX2.sgRNA** (*) symbol at the end indicates that this primer is good to use as probe for sequencing | | | |
|---|---|---|---|
| Gene | Forward (5'-3') | Reverse (5'-3') | Size |
| *Sytabulin* | GCACCGCATGGCTTCTCACC (*) | GGCCCCATCAAAATAAAACCATC | 1.2kb |
| *VEGFA* | TGTGGCGGCCTCCCTTCATCTG (*) | CCCGCTCGCTCGCTCGCTCAC | 887bp |
| *Kinesis* | GCCTGGCACCCTTGACATT | AGCAGGCAGAGCATCCCATCC (*) | 913bp |
| *Stat2* | TTGAGGGGCTGGAGAAAGATAAGT (*) | TGGGGAGCAGAGACAAATAGAGAA | 906bp |
| Chr19 | CACTGCCCACTACCCACTACTAAG (*) | CGGGAGCAATATGGGAAATGGTC | 941bp |

| Set of primers used to screen for off-target of cutting efficiency **RCVRN.sgRNA** (*) symbol at the end indicates that this primer is good to use as probe for sequencing | | | |
|---|---|---|---|
| Gene | Forward (5'-3') | Reverse (5'-3') | Size |
| chr4 | TGTTCCCGGCCATTTGTA (*) | ATCTTGCCAGCATCCATTATCT | 844bp |
| chr2 | AAGCCCACTGGAAAGGTATGAACT (*) | AATGGGAAGGGGACTGAACAAA | 833bp |
| chr14 | AGTTTACGGGAGGGAGGTCAGC (*) | TGGCAGGGAGAAACAGTAGAA | 596bp |
| chr17 | GGGTGGCGGCAGCTTGATAAA (*) | CCCCGAGGATAGCACTGTTGG | 497bp |
| chr17 | GAGCCCCCGGAAGCACAAATACAG (*) | GGCAGGCGTCTCCGTTCTCACAC | 648bp |

| Set of primers used to screen for off-target cutting efficiency of **BRN3b.sgRNA** (*) symbol at the end indicates that this primer is good to use as probe for sequencing | | | |
|---|---|---|---|
| Gene | Forward (5'-3') | Reverse (5'-3') | Size |
| 1100L3.7 | CTTCCCGGCACCAAATCACTCTAC (*) | GCCCCTCCCCTGCTTATCTGG | 1.0kb |
| 45M11.7 | ACCCCTTTTATTCGTGCTCTATTG (*) | AGTCCCGCGTCCTGCTCTC | 1.0kb |
| FAM83F | TGGCCTTTTGCTTTTTCACACC | CACCCCCGGCGTCCTTTACCTG (*) | 854bp |
| RP11-484K9 | CCGTAGGGGGCGAGGAACC (*) | GTGAAGGCGGAAATACAAACAGTC | 691bp |
| RP11-321M21 | GGGGCAAGCTTCTCCACTATTATC | GTTCCATCCTGCGGCTCTTC (*) | 931bp |

Supplemental Table S6: Primers for RT-qPCR

| Gene | Forward (5'-3') | Reverse (5'-3') |
|---|---|---|
| Oct4 | TGTACTCCTCGGTCCCTTTC | TCCAGGTTTTCTTTCCCTAGC |
| NANOG | CAGTCTGGACACTGGCTGAA | CTCGCTGATTAGGCTCCAAC |
| PAX6 | CGGAGTGAATCAGCTCGGTG | CCGCTTATACTGGGCTATTTTGC |
| SIX3 | CCGGAAGAGTTGTCCATGTT | CGACTCGTGTTTGTTGATGG |
| VSX2 | TCATGGCGGAGTATGGGCT | TCCAGCGACTTTTTGTGCATC |
| BRN3a | GGGCAAGAGCCATCCTTTCAA | CTGTTCATCGTGTGGTACGTG |
| BRN3b | CTCGCTCGAAGCCTACTTTG | GACGCGCACCACGTTTTTC |
| RCVRN | CCAGAGCATCTACGCCAAGTT | CCGTCGAGGTTGGAATCGAAG |
| MITF | GACATGCGCTGGAACAAGGGAACC | CCGGGGGACACTGAGGAAAGGAG |
| BEST-1 | AACTGAGCCTACCACACAACA | CGGATTCGACCTCCAAGCC |

Supplemental Table S7: Antibodies used for immunohistochemistry

| Antibodies | Supplier | Species | Type | Dilution | Reference |
|---|---|---|---|---|---|
| VSX2 | Millipore | Sheep | Polyclonal | 1:500 | ab9016 |
| CFP | Abcam | Rabbit | Polyclonal | 1:100 | ab6556 |
| BRN3 | Santa Cruz | Goat | Polyclonal | 1:1000 | sc-6026X |
| MCM2 | Abcam | Rabbit | Polyclonal | 1:1000 | ab4461 |
| Prox-1 | Millipore | Rabbit | Polyclonal | 1:2000 | ab5475 |
| Cralbp | Abcam | Mouse | Monoclonal | 1:500 | ab15051 |
| Ap2-alpha | DSHB | Mouse | Monoclonal | 1:35 | 3B5a |
| Recoverin | Millipore | Rabbit | Polyclonal | 1:500 | ab5585 |
| Oct4 | Abcam | Rabbit | Polyclonal | 1:500 | ab19857 |
| Sox2 | Santa Cruz | Goat | Polyclonal | 1:500 | Sc-17319 |
| Pax6 | Santa Cruz | Mouse | Polyclonal | 1:100 | Sc-32766 |
| Six3 | Santa Cruz | Mouse | Polyclonal | 1:100 | Sc-365519 |
| Rx | Santa Cruz | Mouse | Polyclonal | 1:150 | Sc-271889 |

Supplemental Table S8: RT-qPCR primers for FACS sorted cells

| **Gene** | **Forward (5’- 3’)** | **Reverse (5’ - 3’)** |
|---|---|---|
| Cerulean | AAGCTGACCCTGAAGTTCATCTGC | CTTGTAGTTGCCGTCGTCCTTGAA |
| VSX2 | TCATGGCGGAGTATGGGCT | TCCAGCGACTTTTTGTGCATC |
| mCherry | GATAACATGGCCATCATCAAGGA | CGTGGCCGTTCACGGAG |
| RCVRN | CCAGAGCATCTACGCCAAGTT | CCGTCGAGGTTGGAATCGAAG |
| eGFP | GACCAAAAGATCATGGTGAGC | GAACTTCAGGGTCAGCTTGC |
| BRN3b | CTCGCTCGAAGCCTACTTTG | GACGCGCACCACGTTTTTC |
| GAPDH | CAATGACCCCTTCATTGACC | GACAAGCTTCCCGTTCTCAG |

**Supplemental Figure 1**

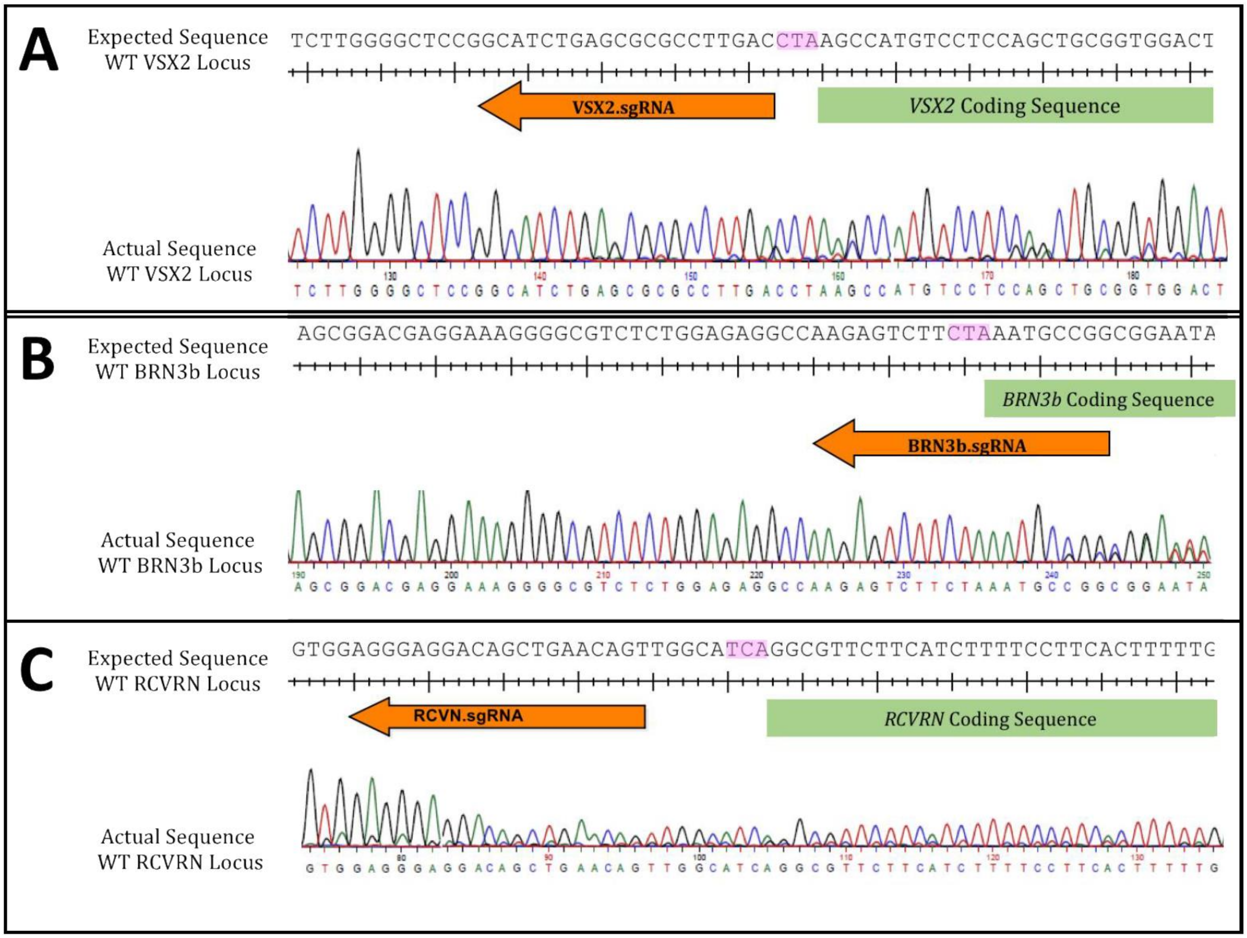

**Supplemental Figure 2**

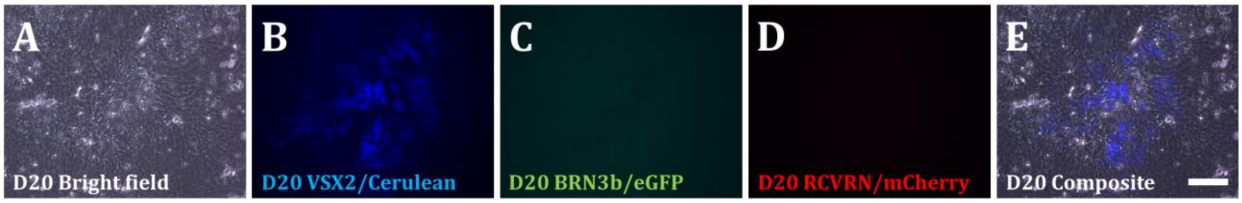

**Supplemental Figure 3**

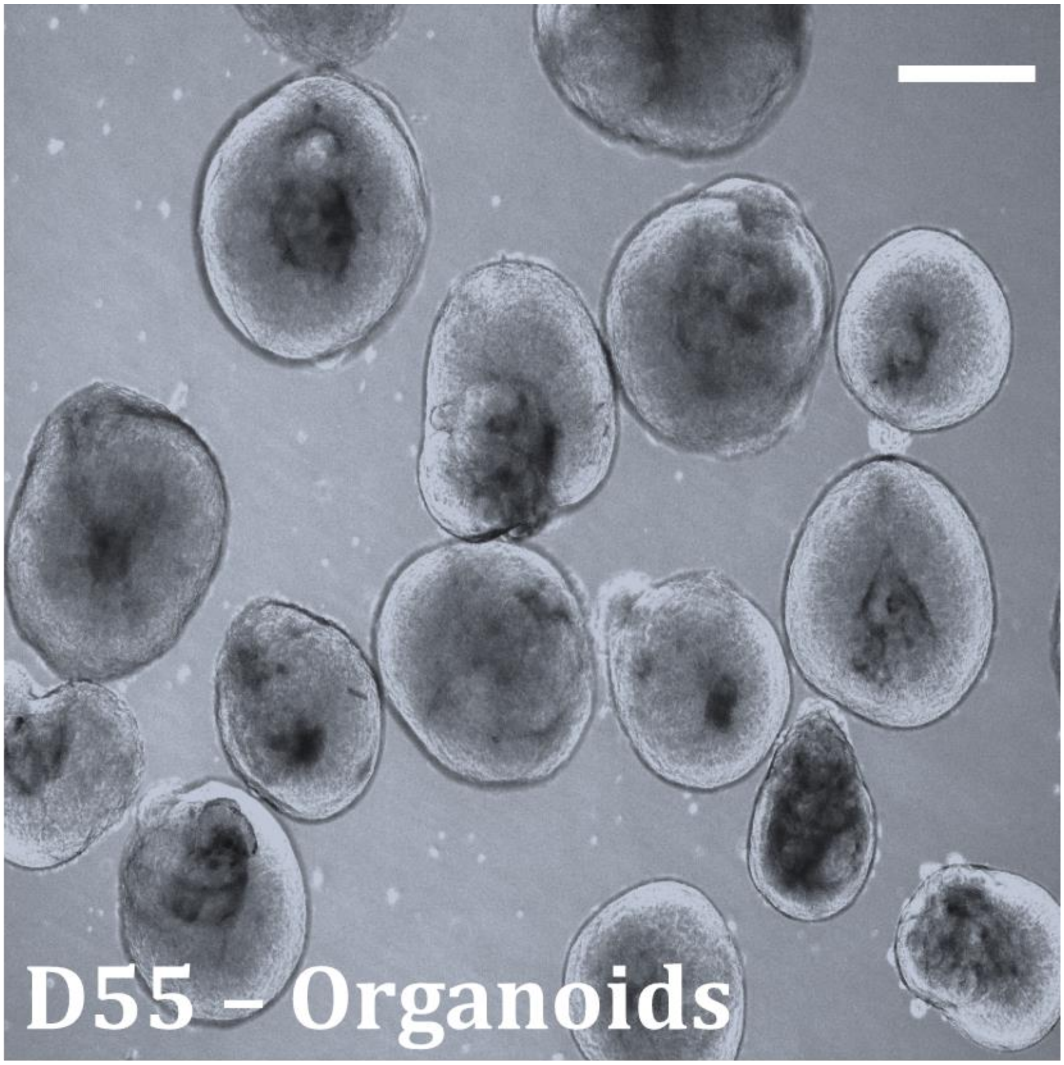

**Supplemental Figure 4**

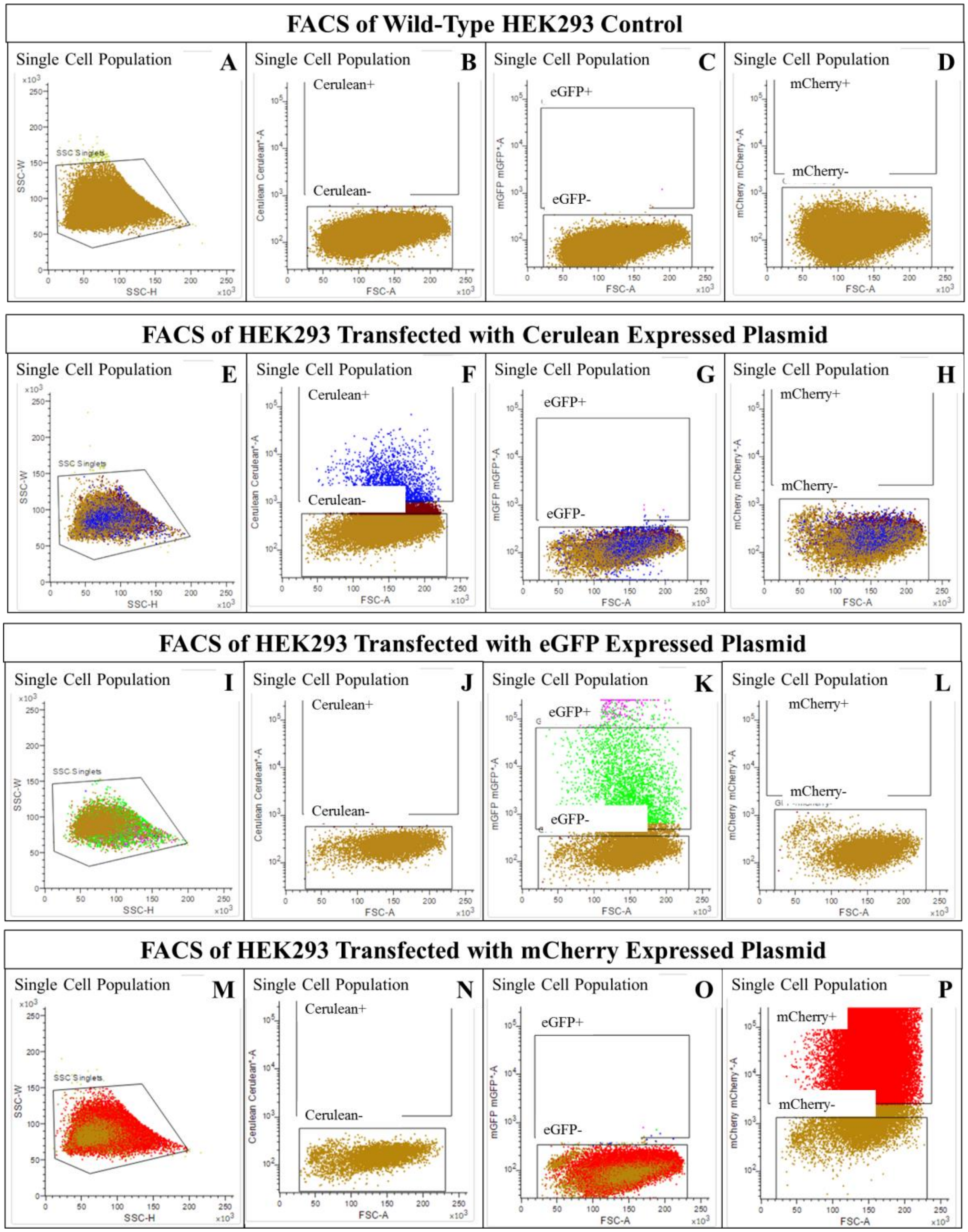

**Supplemental Figure 5**

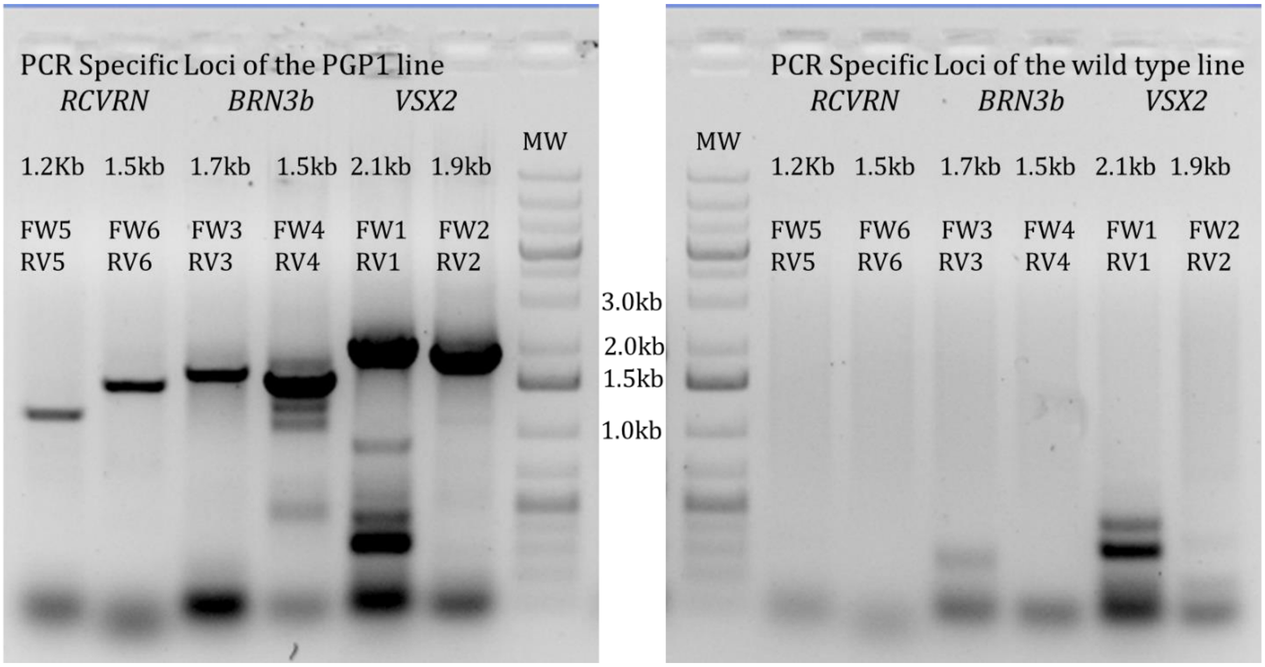

### Supplemental Figure 6

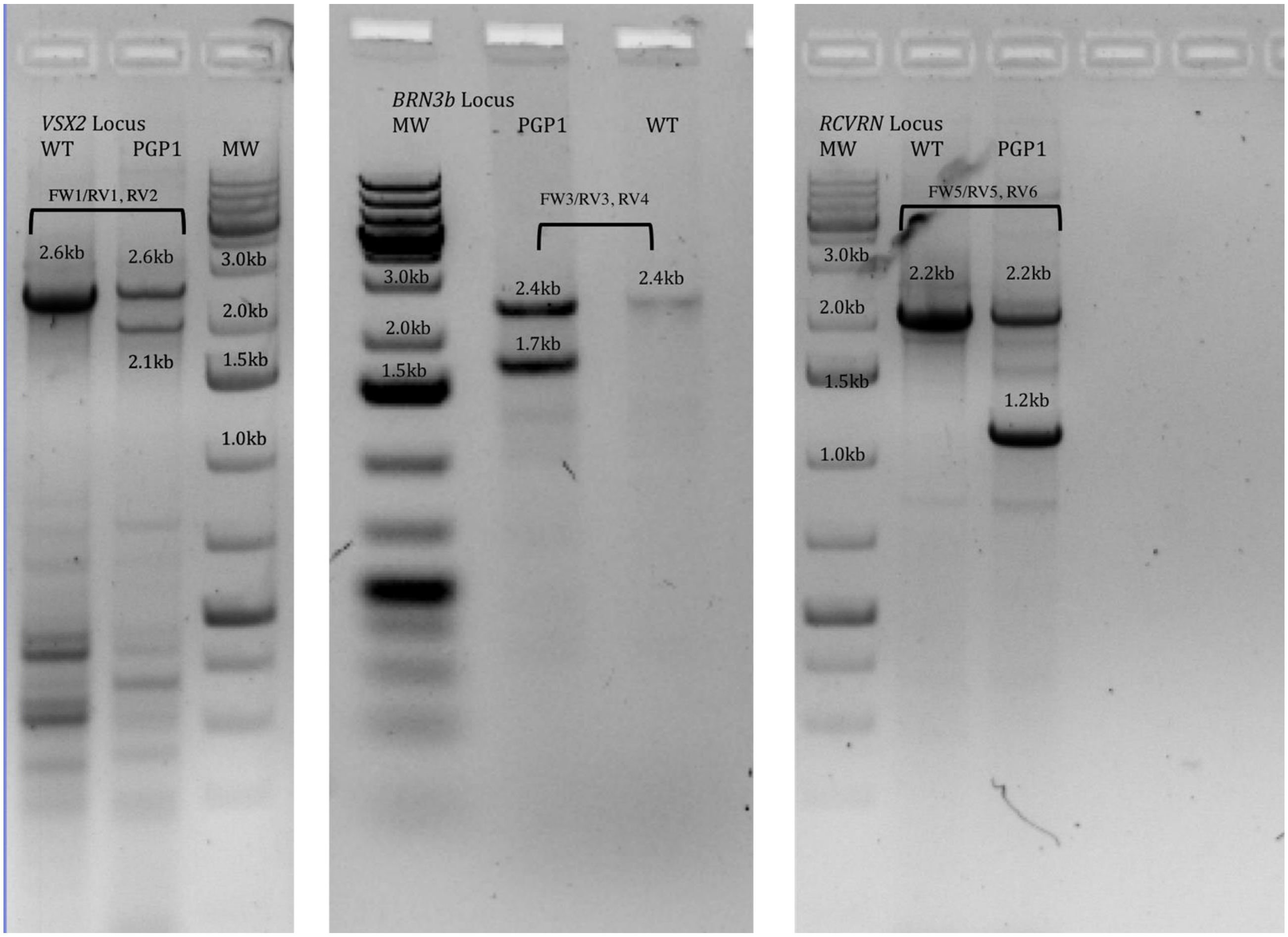

**Supplemental Figure 7**

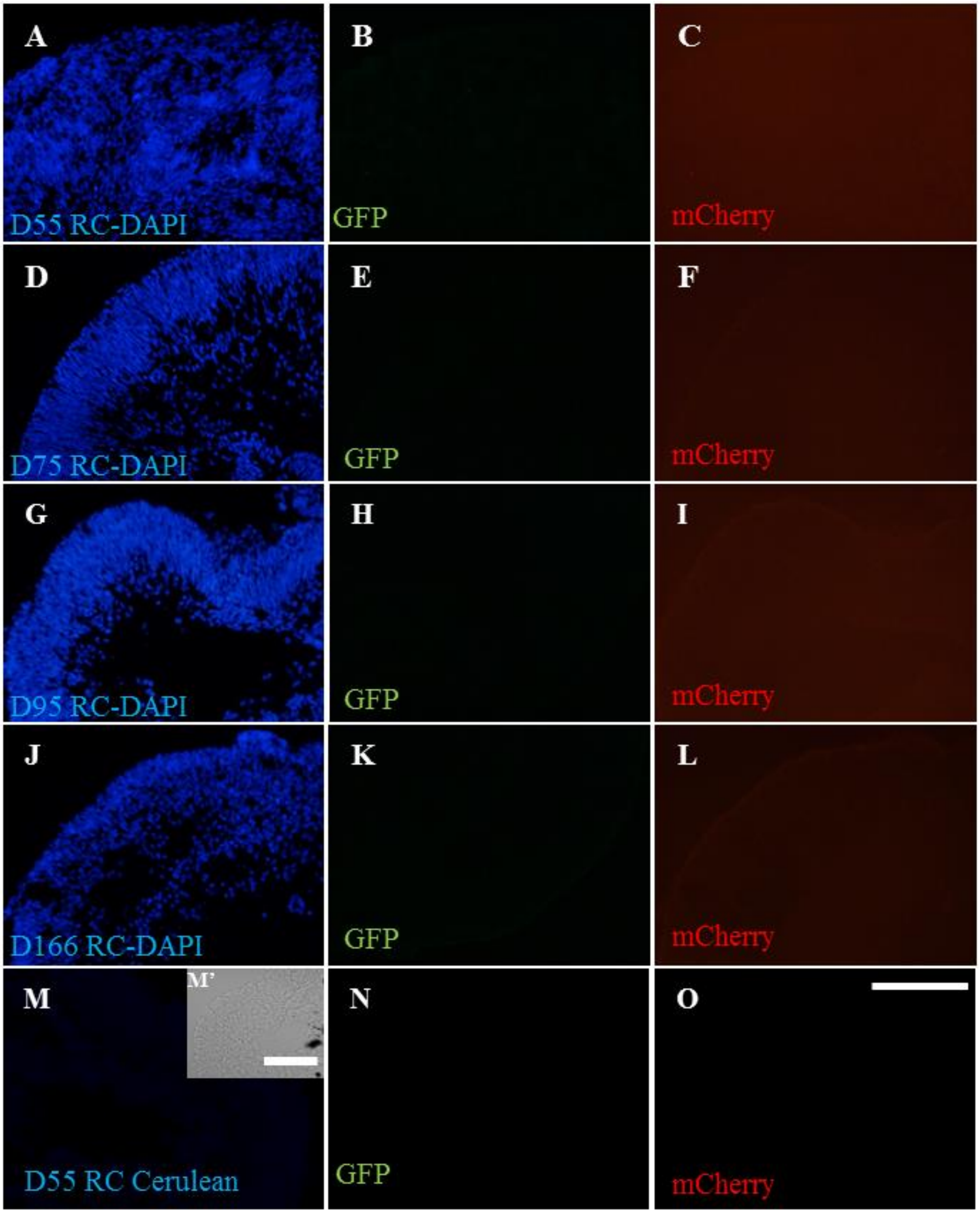

**Supplemental Figure 8**

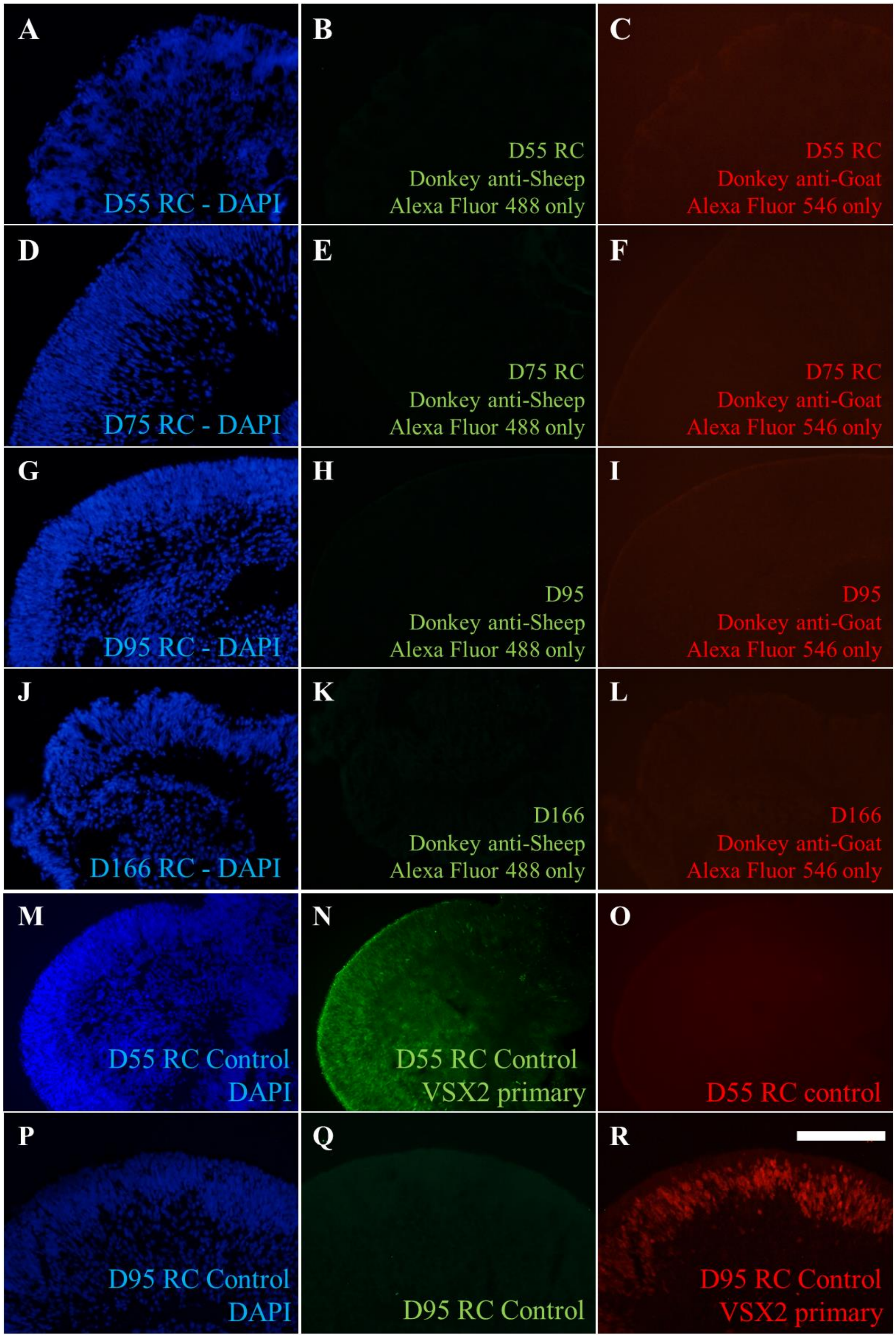

### SUPPLEMENTAL FIGURE LEGENDS

**Supplemental Figure 1: Lack of Indel Mutations in the Non-Targeted Alleles of PGP1.** Sequence analysis of the WT alleles of VSX2 (A), BRN3b (B) and RCVRN (C) failed to detect any Cas9-mediated indel mutations in PGP1. Orange arrows represent the sgRNA sequence, the endogenous stop codons are shaded in pink and coding sequences are represented by green rectangles.

**Supplemental Figure 2**: **Cerulean Positive Retina Progenitors Appear Before eGFP or mCherry Positive Cells During PGP1 Retinal Organoid Differentiation**. After 20 days of differentiation, retinal domains (A) first express Cerulean (blue) (B), but not eGFP (C) or mCherry (D). The composite of the bright field and VSX2/Cerulean (E). Magnification bar 20μm.

**Supplemental Figure 3: Brightfield View of Three Dimension Retinal Organoids at D55 of differentiation.** Free-floating organoids have variable size but all maintain a three dimensional shape with characteristic spherical structure with a distinct thick exterior and hollower interior. Scale bar 40μM

**Supplemental Figure 4: Establishing FACS Gates Using Transiently Transfected HEK293 Cells.** Wild-type HEK293 cells (A-D) or HEK293 cells transiently transfected with expression plasmids for Cerulean (E-H), eGFP (I-L), or mCherry (M-P) were dissociated into single cell populations (A, E, I, M) and Gates were established for each fluorescent protein based on parameters that would lead to capturing the appropriate fluorescent protein expressing cells without capturing any wild-type cells. We confirmed that the captured cells in the Cerulean positive gate (F) the eGFP positive gate (K) and the mCherry positive gate (P) expressed the appropriate fluorescent protein when sorted and cultured.

**Supplemental Figure 5: Original PCR Gels Supporting Figure 1.** The original ethidium bromide stained PCR gels used to support figure 1. PCR reactions using genomic DNA as template with the primers indicated above each lane. The expected band sizes for each targeted allele are shown above each lane. The template DNA for the gel on the left came from the PGP1 clone while the template for the gel on the right was from a wild-type hiPSC clone. MW indicates a DNA size ladder run on each gel.

**Supplemental Figure 6: Original PCR Gels Supporting Figure 2.** The original ethidium bromide stained PCR gel that was cropped for clarity in figure 2. The PGP1 cell line and wild-type hiPSCs (WT) provided the genomic DNA template for a three primer PCR strategy to detect the wild-type and targeted alleles for the *VSX2, BRN3b* and *RCVRN* loci. The primers used for each reaction are indicated above the relevant lanes. MW indicates DNA size ladders run in duplicate on the gel.

**Supplemental Figure 7: Confirmation that Fluorescent Protein Expression in PGP1-Derived Retinal Cup Organoids Does Not Survive Fixation and Frozen Sectioning.** Sections of PGP1 hiPSC-derived retinal organoids were prepared after fixation in 4% paraformaldehyde, overnight incubation in 30% sucrose at 4ºC, and embedding in OCT compound. Organoid sections from D55 (A-C), D75 (D-F), D95 (G-I), and D166 (J-L) of differentiation were visualized following DAPI staining on the blue DAPI filter (A, D, G, J), the green FITC filter for GFP (B, E, H, K) and the red Texas Red filter for mCherry (C, F, I, L). No green or red signals consistent with eGFP or mCherry were detected in organoids of any age. Using an LSM 800 confocal system, an unstained organoid section from D55 was visualized for cerulean expression (Ex.433nm, Em.475) (M), eGFP (Ex.493nm, Em. 517) (N), and mCherry (Ex.577, Em.603) (O). The insert (M') showed the D55 RC section in brightfield. Ex is excitation wavelength and Em is Emission wavelength. Magnification bar for (M') is 50 µm, and 100 µm for all other images.

**Supplemental Figure 8: Additional Controls for PGP1-Derived Retinal Cup Organoids Prepared for Immunohistochemistry.** To ensure the signals from immunofluorescent staining experiments are specific for the intended antigens, the retina cup organoids were stained for DAPI and secondary antibody only (A-L). The staining for DAPI, and the secondary antibodies Donkey anti Sheep Alexa Fluor 488, and Donkey anti Sheep Alexa Fluor 546 were done for sections from organoids at D55 (A-C), D75 (D-F), D95 (G-I), and D166 (J-L) of differentiation. To ensure that the DAPI signal did not represent the VSX2-Cerulean signal, D55 organoid sections were stained with DAPI, and a sheep anti-VSX2 primary antibody and secondary anti sheep Alexa Fluor 488 antibody (M-O). D95 organoid sections were stained with DAPI, and a sheep anti-VSX2 primary antibody and a secondary antisheep Alexa Fluor 546 antibody (P-R). Note that only a subset of the DAPI signals in (M) and (P) are VSX2 positive (N, R). Images in the first column (A, D, G, J, M, P) were photographed with a DAPI filter, while images in the middle (B, E, H, K, N, Q) were photographed with a FITC filter and the last column (C, F, I, L, O, R) were photographed with a Texas Red filter. Magnification Bar. 50µm, applies to all images.

**Supplemental Movie 1: Three-dimensional image of a Day 55 retinal cup Organoid made from the PGP1 cell line.** A portion of a retinal cup organoid made from the PGP1 cell line was imaged at 55 days of differentiation for the expression of Cerulean, eGFP and mCherry to visualize cells expressing *VSX2*, *BRN3b* and *RCVRN*, respectively. At this stage many Ceruelan-expressing cells are found in the organoid. At this stage of development, eGFP-expressing cells exhibit long extensions typical of ganglion cell axons and only a few mCherry-expressing cells appear at this stage. Images taken on an Olympus FV3000 confocal system.
